# Ligand-dependent Enhancer Activation Indirectly Modulates Non-target Promoters in a Chromatin Domain

**DOI:** 10.1101/2024.08.26.609711

**Authors:** Darshika Bohra, Zubairul Islam, Sundarraj Nidharshan, Aprotim Mazumder, Dimple Notani

**Affiliations:** Tata Institute of Fundamental Research Hyderabad, Hyderabad-500046, Telangana, India; National Centre for Biological Sciences, Tata Institute for Fundamental Research, Bangalore-560065, Karnataka, India; Sastra Deemed University, Thanjavur - 613401, Tamil Nadu, India; School of Biotechnology, Amrita Vishwa Vidyapeetham, Kollam - 690525, Kerala, India

## Abstract

Transcription activation of genes by estrogen is driven by enhancers, which are often located within the same Topologically Associating Domain (TAD) as non-targeted promoters. We investigated how acute enhancer-driven activation affects neighbouring non-target genes within the same TAD. Using single-molecule RNA FISH (smFISH), we tracked the transcription of *TFF1* (enhancer-target gene) and *TFF3* (non-target gene) during estrogen stimulation. We observed mutually exclusive expression patterns: TFF1 expression peaked at 1 hour, while TFF3 reached its peak at 3 hours, after TFF1 activation had diminished. Chromatin looping data indicated that the enhancer loops with *TFF1* but not *TFF3*, suggesting that TFF3 upregulation is not due to direct enhancer-promoter interactions. CRISPR deletion of the enhancer, affected TFF1 transcription more acutely than TFF3. 1,6-hexanediol (HD) exposure suggested that the *TFF1* enhancer:promoter undergo a potential ERα-mediated condensate formation, which sequesters the transcriptional machinery and inhibits TFF3 expression. As estrogen signalling fades at 3h, TFF1 expression declines while TFF3 expression increases. Our findings reveal that enhancer-driven activation can indirectly repress neighbouring genes within the same TAD, highlighting a dynamic shift in gene expression as signalling progresses.

## Introduction

Acute transcriptional activation of ligand-induced genes drives downstream signalling response. Similar to development-specific genes, signalling induced genes are also driven by enhancers (Hah et al., 2013; Li et al., 2013; Liu et al., 2014; Uyehara & Apostolou, 2023). Upon binding with ligand-induced transcription factors (TFs), these enhancers loop with their target promoters mostly, in the same TAD (Buecker et al., 2014; Bulger & Groudine, 1999; Chepelev et al., 2012; Furlong & Levine, 2018; Oh et al., 2021; Panigrahi & O’Malley, 2021; Ptashne, 1986; Sanyal et al., 2012; Yan et al., 2018). Enhancer: promoter pairing is thought to be specific and forms the basis of noise-free gene activation of a subset of genes crucial for signalling response (Bojcsuk et al., 2017; H. Chen et al., 2018; Friedman et al., 2024; Galouzis & Furlong, 2022; Zabidi et al., 2015). Due to such specificity, gene transcription occurs in waves of early and late responsive genes (Fowler et al., 2011; Yamamoto & Alberts, 1976). Often, the protein factors translated from early genes regulate the expression of late responsive genes (Dixon et al., 1996; Freter et al., 1996; Herschman, 1991; Williams et al., 1999; Winkles, 1997; Winston & Pledger, 1993). However, it is not known if the genes that are activated early in the signalling time course are spatially related to late activating genes. Further, if the spatial proximity of any gene to an early gene is enough to cause the temporal differential transcription due to sequestration of the transcriptional machinery from late gene to early gene is poorly understood. Transcription factors (TFs), after binding to cognate DNA motifs, recruit RNA polymerase machinery and co-activators for gene activation (Levine & Tjian, 2003; Liu et al., 2014). These machinery are limited in supply, and may become sequestered from other genomic regions (Koşar & Erbaş, 2022). Further, acute activation of genes is linked with phase-separation of TFs, polymerases, mediators and other co-factors/activators, most of which harbour low complexity regions to promote weak protein-protein interactions driving the formation of TF-condensates (Boehning et al., 2018; Boija et al., 2018; Cai et al., 2019; L. Chen et al., 2023; Cho et al., 2018; Chong et al., 2018, 2022; Mann & Notani, 2023; Sabari et al., 2018; Shrinivas et al., 2019; Stortz et al., 2020, 2024). Though the precise stoichiometry of protein molecules in these condensates is unknown, such structures involve multiple molecules of each transcriptional protein. The formation of such phase-separated compartments can potentially act as a sink for transcriptional machinery depriving neighbouring promoters and enhancers that are not part of the compartment. Such sequestration would cause indirect suppression of these spatially proximal genes in the same TAD but not the part of the same condensate.

In order to investigate the effect of an enhancer: promoter pair on a neighbouring gene within the same TAD, we looked at the paradigmatic model of estrogen signalling in mammary epithelial cells, namely MCF7 (Levenson & Jordan, 1997; Masiakowski et al., 1982). E2-signalling can be induced within minutes by treating the cells with estradiol (E2), which causes activation and repression of several genes across the genome. The peak of E2-mediated signalling occurs 40 minutes post-induction and starts to decay by 160 minutes (Hah et al., 2011). This occurs via the binding of estrogen receptor-alpha (ER*α*) on the enhancers (Li et al., 2013). We selected E2-induced E-P pair *TFF1* and its neighbouring gene *TFF3* on Chr21 as discussed below (Chinery et al., 1996). Several studies have used single-cell methods to interrogate E2 responsive genes such as *GREB1*, *MYC* and *TFF1* as a paradigm to understand their transcriptional response upon estrogen stimulation (Fritzsch et al., 2018; Patange et al., 2022; Rodriguez et al., 2019; Stossi et al., 2020). These studies have implicated the cellular states as determinants of transcriptional response and that the long repressive states of TFF1 give rise to expression variability in isogenic lines (Rodriguez et al., 2019).

*TFF1* is found in a topologically associated domain (TAD) along with a few other genes, including *TFF2*, *TFF3*, *TMPRSS3*, and *UBASH3A* (Oh et al., 2021; Quintin et al., 2014; Rao et al., 2014). *TFF1* and *TFF3* is located at 10kb and 60kb from enhancer, respectively (Figure 1A), and this locus has been useful in answering the questions about promoter-enhancer interactions and multi-gene regulation (Oh et al., 2021; Quintin et al., 2014;).

**Figure 1.**
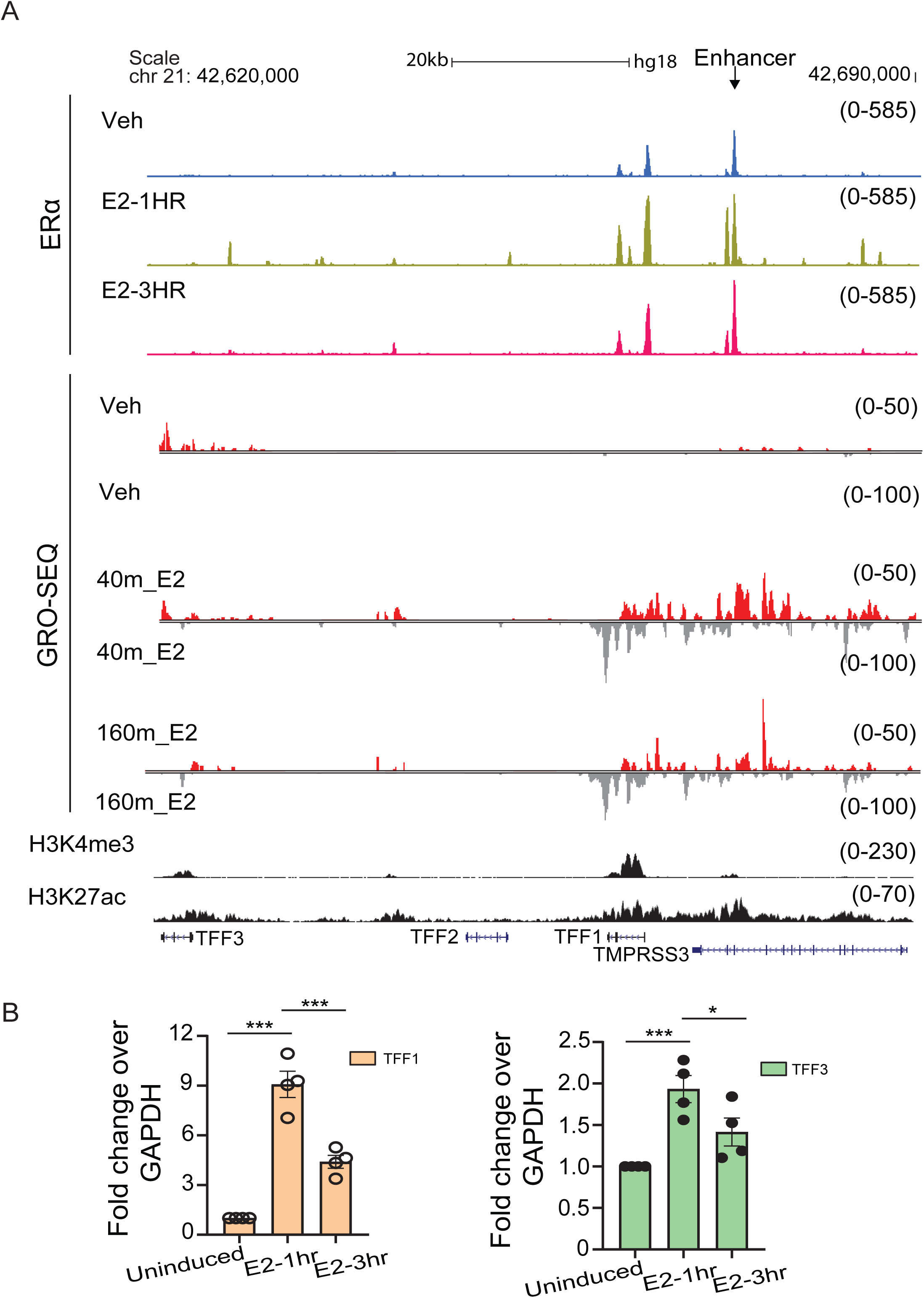
ERα binding and ligand-induced gene expression of *TFF1* and *TFF3* change over the course of estrogen signaling. A. Schematic depicting *TFF1* locus, UCSC genome browser snapshots showing the binding of ERα, H3K27ac status, *H3K4me3* signal, and Gro-seq signal for robustly E2-induced *TFF1* locus. First, second and third ERα ChIP-seq and Gro-seq tracks are from vehicle-treated, E2-1h and E2-3h in WT cells, respectively. B. qRT-PCR showing the changes in expression of *TFF1* and *TFF3* genes during the E2 signaling time course. Error bars denote SEM from four biological replicates. Each dot represents a replicate. p-values were calculated by Student’s two-tailed unpaired t-test, and the significance is represented as: *** denotes p < 0.001,** denotes p < 0.01,* denotes p < 0.05, ns denotes p > 0.05.

We chose to answer these questions using an integrated approach consisting of single molecule RNA FISH (smFISH), genome-wide conformation capture and sequencing (4C-seq), and perturbation of cis-acting regulatory elements by CRISPR. We identified that *TFF1* and *TFF3* genes located within the same TAD show distinct and opposing expression profiles over the course of E2-mediated signalling. In agreement with this, the enhancer also showed increased lopping interaction with the cognate gene promoter during peak expression at 1h of signalling. We identified a potential role of TF-mediated condensates on the up-regulation of TFF1 and down-regulation of TFF3 at the peak of signalling. Hence, we propose that ligand-dependent ERα-protein assemblies can support the expression of the cognate gene in concert with the enhancer that indirectly represses expression of a neighbouring gene in the same TAD.

## Results

### ERα binding and ligand-induced gene expression of *TFF1* and *TFF3* change over the course of estrogen signaling

In order to understand how acute activation of one gene in a TAD affects the transcription of a proximate gene, we chose to study *TFF1* and its neighbouring gene *TFF3* at a 43kb distance within the same TAD. TFF1 expression is linked with an enhancer located 10 kb downstream (Li et al., 2013; Saravanan et al., 2020; Oh et al., 2021) and the distance between *TFF1* enhancer and *TFF3* gene is 53kb (Figure 1A).

17-β-oestradiol exposure leads to ERα binding on regulatory regions, leading to gene activation. Gene transcription of E2-regulated genes was shown to peak at 1h and significantly reduced at 3h due to the rapid degradation of ERα (Hah et al., 2011). The binding of ERα in the genome also follows this temporal kinetics, where it peaks at 1h and reduces at 3h post ligand stimulation (Hah et al., 2011; Li et al., 2013; Liu et al., 2014). At the *TFF1* locus, the *TFF1* enhancer and to some extent, its promoter, were bound by ERα even in the absence of E2 induction, while it increased in strength at these regions and also at various other regions in the locus at 1h of ligand stimulation (Saravanan et al., 2020). Notably, the binding at these regions substantially decreased at 3h. On the other hand, ER*α* did not bind on the *TFF3* promoter throughout the course of signalling (Figure 1- figure supplement 1B). A single peak of ERα was observed approx. 3 kb upstream to *TFF3* gene. We have previously shown that estrogen-induced clustered binding of ER*α* on *TFF1* region is associated with its acute activation post estrogen stimulation (Saravanan et al., 2020). We then tested their expression level by nascent RNA-seq (GRO-seq). We observed a dramatic transcriptional activation of *TFF1* at 1h, which reduced substantially at 3h (Figure 1A). The expression level of *TFF3* was comparatively lower and exhibited mild induction at 1h which increased furthermore at 3h upon signalling (Figure 1- figure supplement 1B). However, qRT-PCR on unspliced TFF3 exhibited upregulation at 1h and down-regulation at 3h (Figure 1B). We reasoned that these differences in TFF3 could be because of its low baseline expression (Clark et al., 2015; Conesa et al., 2016; Sha et al., 2015; Svensson et al., 2017; Tarazona et al., 2011), and lower yet expression of unspliced transcripts. Therefore, in order to test the relative induction of these genes and their co-regulation, we decided to perform single-cell measurements of RNA (Figure 1- figure supplement 1A) that do not rely on PCR amplification and can provide absolute numbers of TFF1 and TFF3 transcripts in a given cell. Towards this, we chose to employ smFISH (Femino et al., 1998; Haimovich & Gerst, 2018; Kwon, 2013; Raj et al., 2008).

### TFF1 and TFF3 exhibit opposite trends during the E2 signalling time-course

MCF7 cells are hypertriploid to hypotetraploid, and each of the three alleles within a nucleus can behave differently, and smFISH is pre-eminently suited to capturing cell- and allele-specific heterogeneity of expression even for low expressing genes, compared to bulk studies that average over cell populations. Briefly, smFISH allows to visualize single RNA molecules in fixed cells using multiple fluorescently labelled oligonucleotide probes targeted to the RNA of interest and therefore, it can be used to image the transcription and the localization of multiple gene transcripts at same and different time points after signalling (Femino et al., 1998; Raj et al., 2008).

We designed the smFISH probes targeting the intronic region of TFF1 and TFF3 to measure their nascent transcripts (referred to as InTFF1 and InTFF3 hereon) whereas the exonic probes were used to primarily measure mature mRNA transcripts (referred to as ExTFF1 and ExTFF3 hereon). Intronic probes are particularly suited for investigating nascent transcriptional status at the time of fixation, while probes against mature mRNA reflects on more steady-state levels of functional mRNA due to finite mRNA lifetimes (Skinner et al., 2016). Additionally, the intronic probes can help determine the localisation and number of alleles that are transcribing at a given time thus allowing for additional interpretations regarding the expression of multiple genes (Skinner et al., 2016).

Representative images from the smFISH experiment probing InTFF1 and InTFF3 are depicted (Figure 2A). The smaller foci represent individual intronic/nascent transcripts while the larger foci represent the site of transcription (Raj et al., 2006; Zenklusen et al., 2008). Since MCF7 cells are hypertriploid in nature, we expected to see 1-3 large foci per cell representing sites of transcription. Additionally, we observed many individual transcripts labelled by intronic probes (Figure 2A, D) within the same nuclei (probes against mature RNA are usually more cytoplasmic). This is suggestive of the fact that the transcripts undergo non-co-transcriptional splicing as the transcripts labelled by intronic probes are localised away from the site of transcription. This is not surprising as several reports have shown that nascent transcripts can undergo splicing well after transcription (Coulon et al., 2014; Drexler et al., 2020; Khodor et al., 2012).

**Figure 2.**
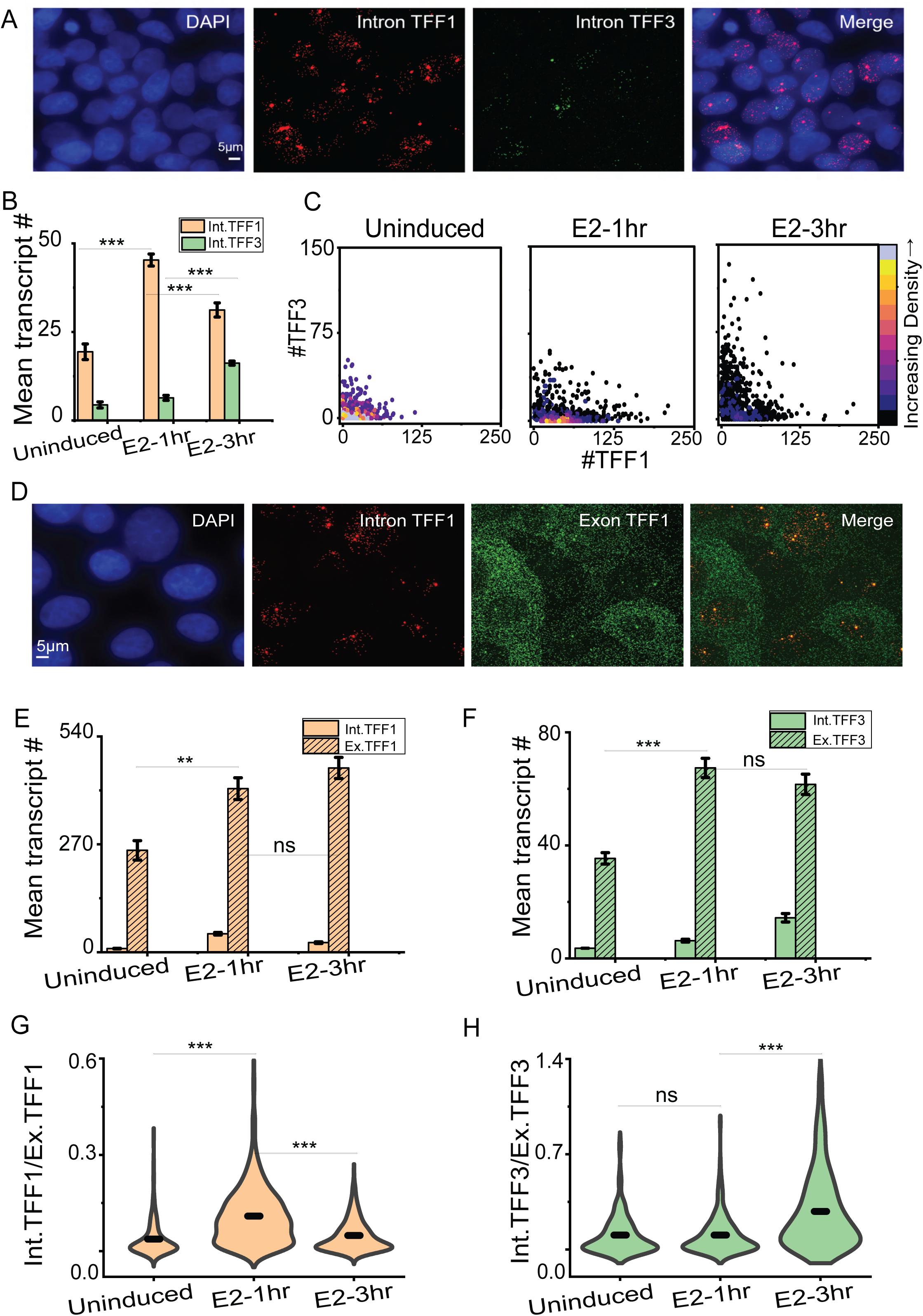
*TFF1* and *TFF3* expressions show opposite trends during the E2 signaling time-course. A. 60X Representative images from single molecule RNA FISH experiment showing transcripts for TFF1 and TFF3. The probe was designed against the unspliced RNA containing the intronic region. The scale bar is 5 microns. B. The mean RNA numbers are depicted. These are counted using an in-house MATLAB code which uses the DAPI-stained nuclei as the mask to count the RNA present in the nucleus. The graph shows the mean of means from three different repeats of the experiment, and error bars denote SEM (n= 665, N=3). p-values were calculated by Student’s two-tailed unpaired t-test, and the significance is represented as: *** denotes p < 0.001,** denotes p < 0.01,* denotes p < 0.05, ns denotes p > 0.05. C. Scatter plots showing the distribution of InTFF1 and InTFF3 on a cell-by-cell basis (n= 665, N=3). The absolute RNA numbers are combined from three different repeats. Density plots have been used to clearly visualize overlapping data points. D. 60X Representative images from single molecule RNA FISH experiment showing transcripts for InTFF1 and ExTFF1. Scale bar is 5 micrometers. E. The mean RNA numbers for InTFF1 and ExTFF1 are depicted. Separate probes were used to target unspliced (InTFF1) and mature (ExTFF1) RNA. These are counted using an in-house MATLAB code which uses the DAPI-stained nuclei as the mask to count the intronic RNA present in the nucleus and a free-drawn region to designate the cell to count the exonic RNA present in the nucleus as well as the cytoplasm. The graph shows the mean of means from three different repeats of the experiment, and error bars denote SEM (n>360, N=3). p-values were calculated by Student’s two-tailed unpaired t-test, and the significance is represented as: *** denotes p < 0.001,** denotes p < 0.01,* denotes p < 0.05, ns denotes p > 0.05. F. The mean RNA numbers for InTFF3 and ExTFF3 are depicted. Separate probes were used to target unspliced (InTFF3) and mature (ExTFF3) RNA. The graph shows the mean of means from three different repeats of the experiment, and error bars denote SEM (n>210, N=3). p-values were calculated by Student’s two-tailed unpaired t-test, and the significance is represented as: *** denotes p < 0.001,** denotes p < 0.01,* denotes p < 0.05, ns denotes p > 0.05. G. Violin plots showing the ratio of intronic to exonic TFF1 counts are depicted. The graph shows the distribution of ratios combined from three different repeats (n>360, N=3). p-values were calculated by the Mann-Whitney test, and the significance is represented as: *** denotes p < 0.001,** denotes p < 0.01,* denotes p < 0.05, ns denotes p > 0.05. H. Violin plots showing the ratio of intronic to exonic TFF3 counts are depicted. The graph shows the distribution of ratios combined from three different repeats (n>210, N=3). p- values were calculated by the Mann-Whitney test, and the significance is represented as: *** denotes p < 0.001,** denotes p < 0.01,* denotes p < 0.05, ns denotes p > 0.05.

We quantified the number of such transcripts per nuclei as a proxy for ongoing transcription. At 1h post E2 induction, TFF1 mean transcript counts increased significantly compared to uninduced, whereas, the increment was less pronounced for TFF3 transcripts. In contrast, the mean transcript counts for TFF3 increased significantly at 3h post induction while TFF1 transcription showed a decrease at 3h compared to 1h (Figure 2B). We observed an increase in the number of transcribing sites per cell for both TFF1 and TFF3, as well as in the number of cells showing transcripts compared to the untreated condition (Figure 2- figure supplement 1A-B). Further, TFF1 transcripts spots were more spread out in the nucleus that harboured multiple transcribing sites (Figure 2-figure supplement 1C-D). To check the transcriptional status of *TFF1* and *TFF3* in the same cell, we plotted the transcript counts from individual cells (Figure 2C). The data suggested that the transcript counts for TFF1 increased at 1h and cells that showed high counts for TFF1 were likely to have low counts for TFF3 and vice-versa. But the RNA counts for TFF1 decreased at 3h and increased for TFF3. Overall, it was evident that the transcriptional profile for these two genes located in the same TAD were negatively correlated as they peaked at different time points. To further confirm this, smFISH using probes targeting both the intronic and exonic transcripts in the same experiment was conducted. Intronic probes represent active transcription, while exonic probes show accumulated mature RNA from past transcriptional events even in the absence of active transcription (Skinner et al., 2016). Representative images from the smFISH experiment probing InTFF1/ExTFF1 and InTFF3/ExTFF3 are depicted (Figure 2D, Figure 2- figure supplement 1E). InTFF1 spots overlapped with ExTFF1 in the nucleus (Figure 2D, Figure 2- figure supplement 1F) and InTFF1 spots were restricted to the nucleus, whereas ExTFF1 were both in the nucleus and the cytoplasm as expected, indicating the specificity of smFISH probes (Figure 2D, Figure 2 - figure supplement 1E). We observed that the exonic transcript counts for both TFF1 and TFF3 increased at 1h compared to uninduced (Figure 2E, F). Strikingly, at 3h, the exonic transcripts for TFF1 continued to increase even while the intronic transcript counts reduced, though the fold increase (1.12 ±0.02) was less compared to that between uninduced and 1h (1.63 ± 0.28), indicating a plateauing of steady state levels. Exonic transcripts for TFF3 at 3h remained comparable to 1h while the intronic counts increased suggesting, TFF1 transcription reduces at 3h, whereas TFF3 expression increases. We reasoned that the ratio of intronic transcript number (InTFF) to exonic transcripts number (ExTFF) should represent the status of transcription as an increase in the number of intronic transcript due to expression would result in a higher ratio while also taking into consideration the number of mature transcripts. As expected, the ratio of intronic transcripts to exonic transcripts also showed that transcription is active at 1h for *TFF1* as the ratio is higher compared to uninduced and 3h (Figure 2G, Figure 2- figure supplement 2A). Contrastingly, the ratio was highest at 3h for *TFF3* suggesting that active transcription takes place much after the peak of E2-mediated signalling and maximal TFF1 expression (Figure 2H, Figure 2- figure supplement 2B). Indeed, a very recent study has shown that such post-transcriptional splicing can occur for genes and intron dispersal can be expected more commonly for some highly expressed genes (Coté et al., 2023). Therefore, the observation of intronic signal away from the site of transcription as we see for TFF1 and TFF3 is not unexpected (Figure 2A, D, Figure 2-figure supplement 1).

### *TFF1* enhancer does not change target promoters during signalling time-course

In order to identify the molecular players behind the differential expression of these two genes that are in the same TAD, we asked if the ER*α*-bound enhancer downstream to the *TFF1* gene loops with *TFF1* at 1h, and with *TFF3* at 3h. This enhancer acutely activates *TFF1* at 1h post estrogen stimulation (Saravanan et al., 2020; Ho et al., 2021). We interrogated the looping using enhancer as a viewpoint by 4C-seq in uninduced, 1h and 3h post E2 stimulation. We observed robust interactions between enhancer and *TFF1* promoter at 1h post induction, which reduced at 3h. On the other hand, its interaction with the *TFF3* promoter exhibited very low counts in uninduced as well as E2-induced conditions at both time points and in both replicates (Figure 3A, Figure 3-figure supplement 1). This suggests that the looping of enhancer potentially induces the expression of *TFF1* gene at 1h, and loss of interactions at 3h results in weak *TFF1* transcription. However, the lack of interactions between enhancer and *TFF3* did not explain the gain of *TFF3* expression at 3h post ligand stimulation.

**Figure 3.**
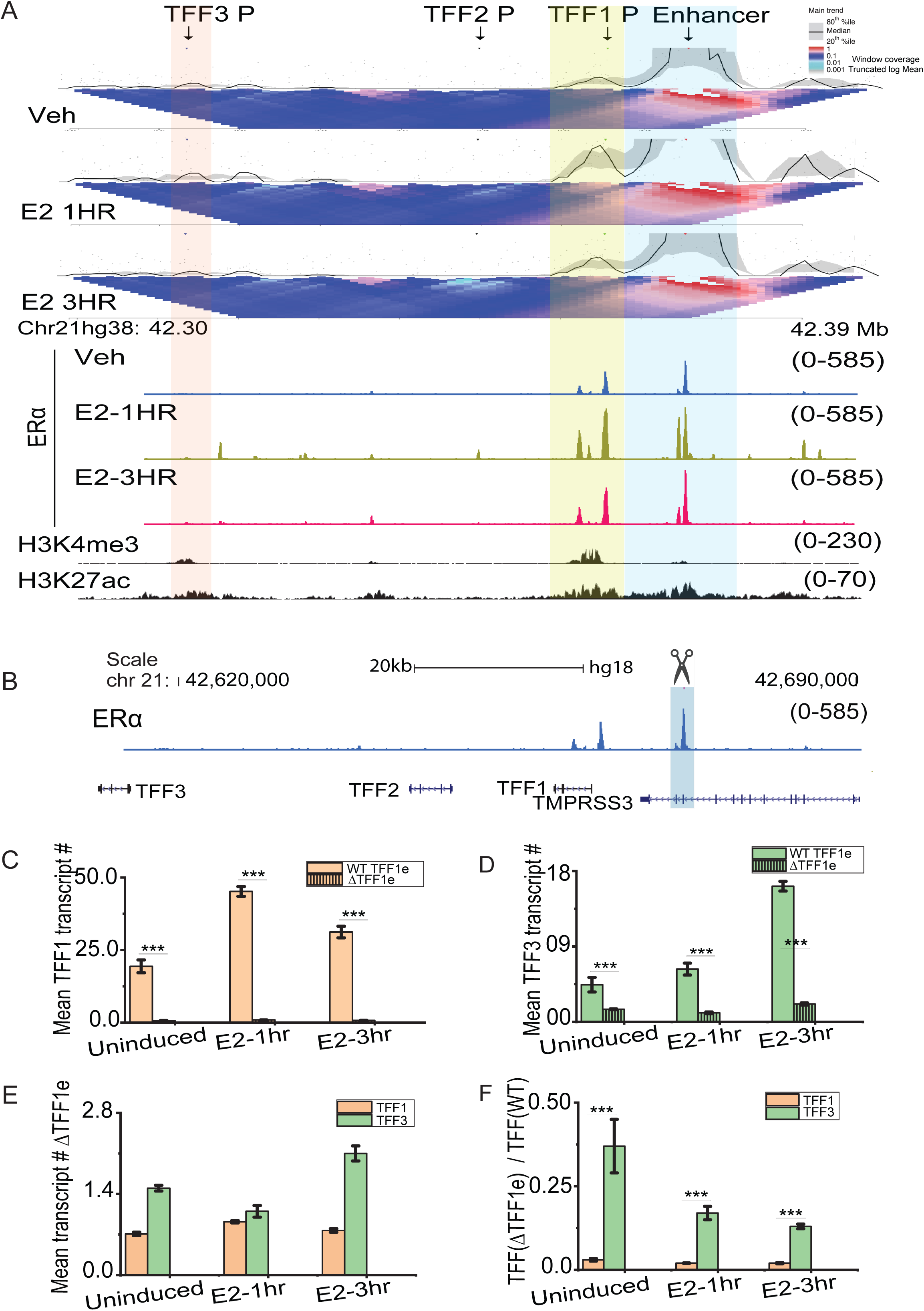
Enhancer looping does not account for the differential expression of *TFF1* and *TFF3* genes. A. 4C-seq plot at *TFF1* enhancer viewpoint, the interaction with the promoter is highlighted in yellow. The plot is overlaid with H3K27ac, ERα ChIP signal, and gene annotations. B. Genome browser snapshot of *TFF1* region depicting ERα binding in WT lines. The first, second and third ERα ChIP-seq tracks are from WT cells that are vehicle-treated, E2-1h, and E2-3h, respectively. Blue highlighted regions represent the Δ*TFF1e* region. C. The mean RNA numbers for InTFF1 in WT (unshaded) and Δ*TFF1e* (shaded) MCF7 cells are depicted. The mean of means are shown, and error bars denote SEM from three repeats (n>650, N=3 for each WT and delete line). p-values were calculated by Student’s two-tailed unpaired t-test, and the significance is represented as: *** denotes p < 0.001,** denotes p < 0.01,* denotes p < 0.05, ns denotes p > 0.05. D. The mean RNA numbers for InTFF3 in WT (unshaded) and Δ*TFF1e* (shaded) MCF7 cells are depicted. The mean of means are shown, and error bars denote SEM from three repeats (n>650, N=3 for each WT and delete line). P-values were calculated by Student’s two-tailed unpaired t-test, and the significance is represented as: *** denotes p < 0.001,** denotes p < 0.01,* denotes p < 0.05, ns denotes p > 0.05. E. The mean RNA numbers for InTFF1 and InTFF3 in Δ*TFF1e* MCF7 cells are depicted. The mean of means are shown, and error bars denote SEM from three repeats (n>880, N=3). p-values were calculated by Student’s two-tailed unpaired t-test, and the significance is represented as: *** denotes p < 0.001,** denotes p < 0.01,* denotes p < 0.05, ns denotes p > 0.05. F. Ratio of InTFF in WT MCF7 to InTFF in Δ*TFF1e* MCF7 are depicted. The ratio was obtained by dividing the absolute RNA counts of the WT line by delete lines performed on different days but in the same order (replicate one of WT divided by replicate one of ΔTFF1e). The mean of means are shown, and error bars denote SEM from three repeats (n>650, N=3 for each WT and delete line). p-values were calculated by Student’s two-tailed unpaired t-test, and the significance is represented as: *** denotes p < 0.001,** denotes p < 0.01,* denotes p < 0.05, ns denotes p > 0.05.

This could mean that enhancer interaction is critical for *TFF1* expression, but is less important for *TFF3* expression. Thus, *TFF1* expression should be affected more severely upon deletion of the enhancer than *TFF3*. To test this, we investigated the expression of *TFF1* and *TFF3* in MCF-7 cells where the enhancer downstream to *TFF1* was homozygously deleted using CRISPR-Cas9 (referred to as the Δ*TFF1*e from hereon) (Figure 3B) (Saravanan et al., 2021). Using smFISH, we quantified the intronic transcripts for TFF1 and TFF3 in the Δ*TFF1*e cells, compared to WT cells. We observed that the mean number of TFF1 transcripts was reduced drastically in the Δ*TFF1*e compared to the WT (Figure 3C). The reduction was far more substantial (49.23 ±2.6 at 1hr and 40.75 ±7.8 at 3h) for TFF1 than TFF3 (6.03 ±1.8 at 1hr and 7.6 ±0.7 at 3h) (Figure 3D). The absolute transcript counts for TFF1 and TFF3 in the Δ*TFF1*e cells have also been shown for clearer visualization (Figure 3E), as these are obscured when compared to WT. To get a sense of fold changes, we took a ratio of mean transcript counts in WT cells to Δ*TFF1*e cells at each time point. We observed that the drop in gene transcripts between WT and Δ*TFF1*e were several folds higher for TFF1 compared to TFF3 (Figure 3F). This suggests that enhancer deletion has a more robust impact on the transcription of TFF1 compared to TFF3. Nonetheless, *TFF3* was also affected even though it does not loop with the enhancer. This is in accordance with the 4C-seq data where we observed prominent looping between the enhancer and *TFF1* but less so with *TFF3*. These results suggest that the enhancer plays a more important role in the expression of the primary gene while it has less impact on a gene located more distally but within the same TAD during the course of signalling.

### Levels of ERα in the nucleus dictate the extent of *TFF1* and *TFF3* inductions

After ruling out enhancer looping as the determinant of differential expression, we looked for other candidates that could regulate the differential gene expression. As discussed above, globally, E2-mediated gene expression is known to peak at 1h after stimulation and drop significantly by 3h (Hah et al., 2011). Upon ligand stimulation, ER*α* translocates into the nucleus, increasing mean ERα intensity in the nucleus which is high at 1h and then decreases significantly by 3h due to degradation. These changes in intensities have been captured using immunofluorescence for ERα (Saravanan et al., 2020). We tested if the intensities of ERα in individual cells correlate with the expression of TFF1 and TFF3 at 1h and 3h of E2 signalling. Towards this, we combined smFISH with immunofluorescence for ERα. To improve contrast for the ERα signal, we additionally performed a chromatin retention assay to get rid of any chromatin unbound ERα (Figure 4A). The representative images show that the cells with very high levels of nuclear ERα (blue circle) exhibited low counts of both TFF1 and TFF3. In contrast, the cells with medium levels of ERα (red circle) possessed higher TFF1 than TFF3. Similarly, the cells with the lower levels of ERα (grey circle) showed higher TFF3 expression as compared to TFF1 (Figure 4A). Histograms depicting the ERα mean intensities across individual cells, showed that the nuclear level of ERα increases post 1h of induction and then goes down at 3h (Figure 4B, C), similar to TFF1 expression (Figure 4D).

**Figure 4.**
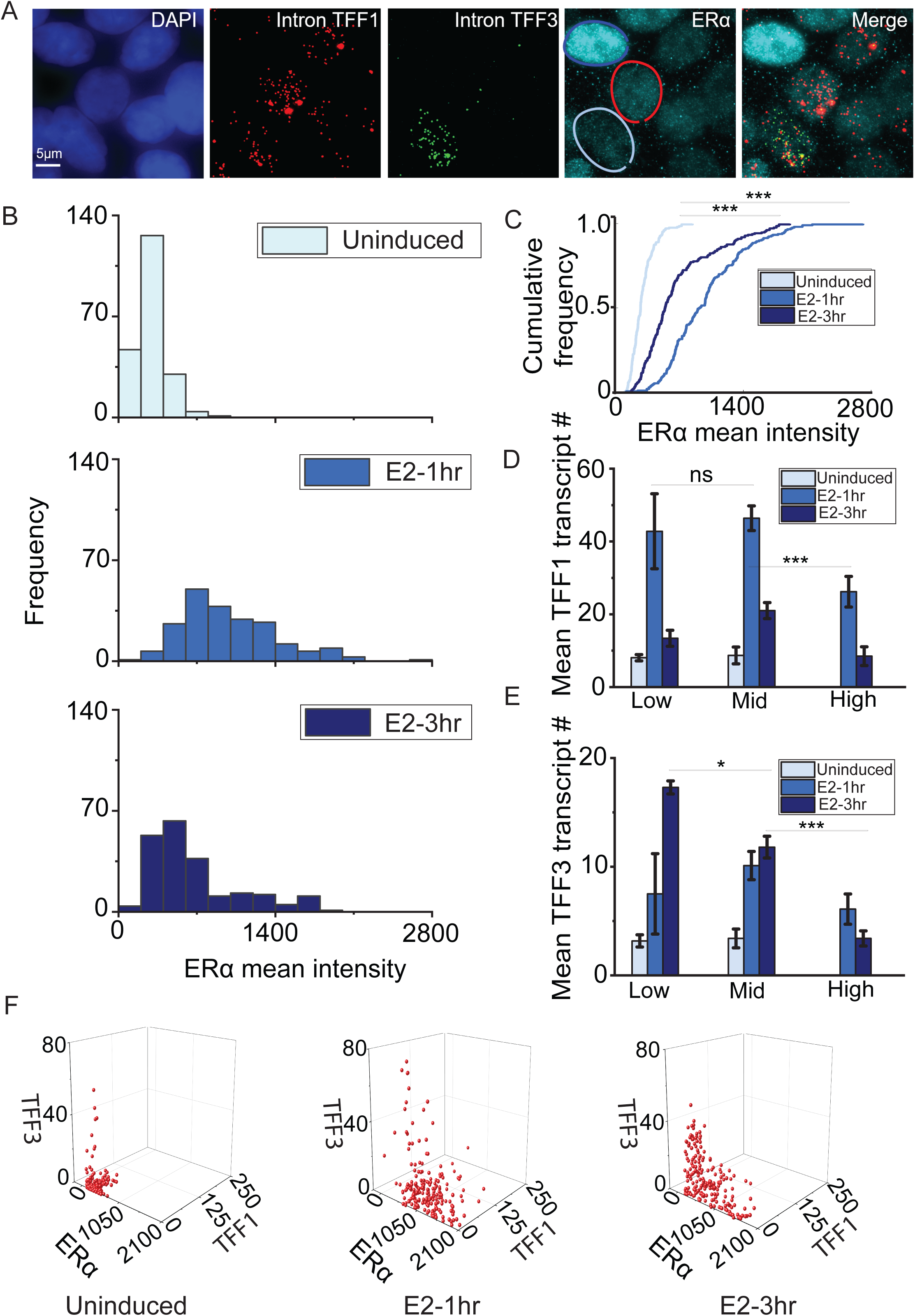
Nuclear levels of ERα dictate the extent of TFF1 and TFF3 expression. A. Representative images showing smFISH for InTFF1 and InTFF3 in combination with immunofluorescence for ERα (along with chromatin retention assay). The red circle denotes a cell with high ERα and low TFF1 and TFF3, the blue circle denotes a cell with medium ERα, high TFF1, and low TFF3 while the green circle denotes a cell with low ERα and high TFF3. The scale bar is 5 micrometres. B. Histogram representing the distribution of ERα mean intensities in cells under uninduced, E2-1hr, and E2-3hr conditions (n=210). Intensities at 1hr are the highest while they shift to the left at 3hr and are lowest in uninduced cells. This is plotted from one experimental repeat out of three repeats as ERα intensities will vary from one immunofluorescence experiment to another. C. Cumulative histogram representing the distribution of ERα mean intensities in cells under uninduced, E2-1hr, and E2-3hr conditions. p-values were calculated by the Mann-Whitney test and the significance is represented as : *** denotes p < 0.001,** denotes p < 0.01,* denotes p < 0.05, ns denotes p > 0.05. D. ERα intensities were sorted into three categories namely low (intensities between 0-450 A.U.) mid (intensities between 450-1200 A.U.) and high (intensities between 1200-2100 A.U.). The mean and SEM of transcript count for InTFF1 in the three categories under uninduced, E2-1hr, and E2-3hr was plotted. Low and mid categories show the highest TFF1 mean. p-values were calculated by the Mann-Whitney test and the significance is represented as : *** denotes p < 0.001,** denotes p < 0.01,* denotes p < 0.05, ns denotes p > 0.05. This is plotted from one experimental repeat out of three repeats as ERα intensities will vary from one immunofluorescence experiment to another. E. ERα intensities were sorted into three categories namely low (intensities between 0-450 A.U.) mid (intensities between 450-1200 A.U.) and high (intensities between 1200-2100 A.U.). The mean and SEM of transcript count for InTFF3 in the three categories under uninduced, E2-1hr, and E2-3hr was plotted. Low category shows the highest TFF3 mean. p-values were calculated by the Mann-Whitney test and the significance is represented as : *** denotes p < 0.001,** denotes p < 0.01,* denotes p < 0.05, ns denotes p > 0.05. This is plotted from one experimental repeat out of three repeats as ERα intensities will vary from one immunofluorescence experiment to another. F. 3D plot representing the distribution of ERα, InTFF1, and InTFF3 on a cell-by-cell basis shows that cells with lower levels of ERα show higher counts for InTFF3. This is plotted from one experimental repeat out of three repeats as ERα intensities will vary from one immunofluorescence experiment to another.

To further corroborate this, we parsed the ERα population cells into 3 categories, namely low, medium, and high within the cells imaged at 1 and 3h. We plotted the mean counts of TFF1 in each of these bins at different time points and observed that the mean count was higher in the mid-category (Figure 4D). While for TFF3, the mean count was significantly higher in the low bin (Figure 4E). The transcript counts for TFF1 and TFF3 against ERα intensities on a cell-by-cell basis (Figure 4F), also showed this feature where very high levels of ERα in the nucleus were not conducive to the expression of either gene (Figure 4F). To test if ER*α* had a causal role in intensity based expression of TFF1 and TFF3, we increased the levels of ER*α* by overexpression of ERα-GFP (Figure 4- figure supplement 1A). The representative images from the smFISH experiment in cells overexpressing ERα-GFP confirmed that *TFF1* and *TFF3* were downregulated in transfected cells while these genes were not perturbed in non-transfected cells in the neighbourhood (Figure 4- figure supplement 1A, C). Meanwhile, the cells overexpressing ERα-GFP do not show any impairment in the expression of GAPDH (housekeeping gene) (Figure 4-figure supplement 1A, B). As another control, we transfected the cells with EGFP-C1 (same backbone as ERα-GFP construct) and observed no effect on TFF1 or TFF3 (Figure 4- figure supplement 1D, E). The data suggest that loss of TFF1 and TFF3 expression upon ER*α* overexpression was not a general effect of transfection stress but rather specific to ERα overexpression. These results indicate that high nuclear levels of ERα can be detrimental to the expression of genes it regulates. This could be due to the widespread condensate formation, which in turn could sequester the transcriptional protein complexes and competitively abrogate transcription across multiple loci. Thus, the global level of ERα in the nucleus can predict the transcriptional status of specific genes. The binding of ER*α* at 1 and 3h is proportional to its nuclear levels (Figure 3A), suggesting its overexpression would lead to more binding in the genome, which is detrimental to gene expression.

### 1,6 HD exposure down-regulates *TFF1* but supports *TFF3* expression

The data obtained from combined smFISH and ERα immunofluorescence indicates that ERα could be a determining factor in controlling the differential gene expression of TFF1 and TFF3. Existing literature indicates that ERα-mediated condensate plays a role in E2-induced gene expression (Boija et al., 2018; Nair et al, 2019; Sabari et al., 2018; Saravanan et al., 2020). It led us to hypothesize that ERα-mediated condensate at the *TFF1* locus could be sequestering all the factors required for active transcription thus preventing activation of the *TFF3* locus at the active phase of E2 signalling. To test this, we treated the cells with 3% 1,6-Hexanediol for 5 minutes, which is known to disrupt LLPS (Gamliel et al., 2022; Kroschwald et al., 2017). Following the treatment, we performed smFISH to look at the transcription of TFF1 and TFF3 in the same cell and observed a dramatic reduction in TFF1 transcripts, whereas a statistically significant increase in TFF3 was noted (Figure 5A). Since ER*α* forms condensate only after estrogen stimulation, 1,6-Hexanediol had no effect on TFF1 in the absence of E2 signalling (Figure 5- figure supplement 1). Together, these results suggest that the functional loss of *TFF1* promoter transcription potentially due to the dissolution of ERα condensate allowed the *TFF3* promoter in the neighbourhood to gain access to transcriptional machinery, leading to its upregulation. In order to test the generality of this observation, beyond the *TFF1* and *TFF3* locus, we divided E2-responsive genes into three categories, low, moderate, and high, based on their expression upon E2-40m stimulation. We observed a significant increase at 40m and down-regulation at 160m when the binding of ERα is reduced in the genome (Figure 5B). Next, we tested the expression of their nearby upregulated genes at 160m compared to 40m post E2 stimulation. The nearby genes are within a distance of 1Mb from the E2-responsive genes, which is approximately the average size of a TAD. Indeed, we observed a significant upregulation of nearby genes at 160m when the expression of highly induced genes dropped. Additionally, the expression of these neighbouring genes was reduced at the peak of signalling (Figure 5B), showing that at 40m of signalling, the acute activation of primary genes and sequestration of transcription machinery by these genes leads to the loss of expression of nearby genes.

**Figure 5.**
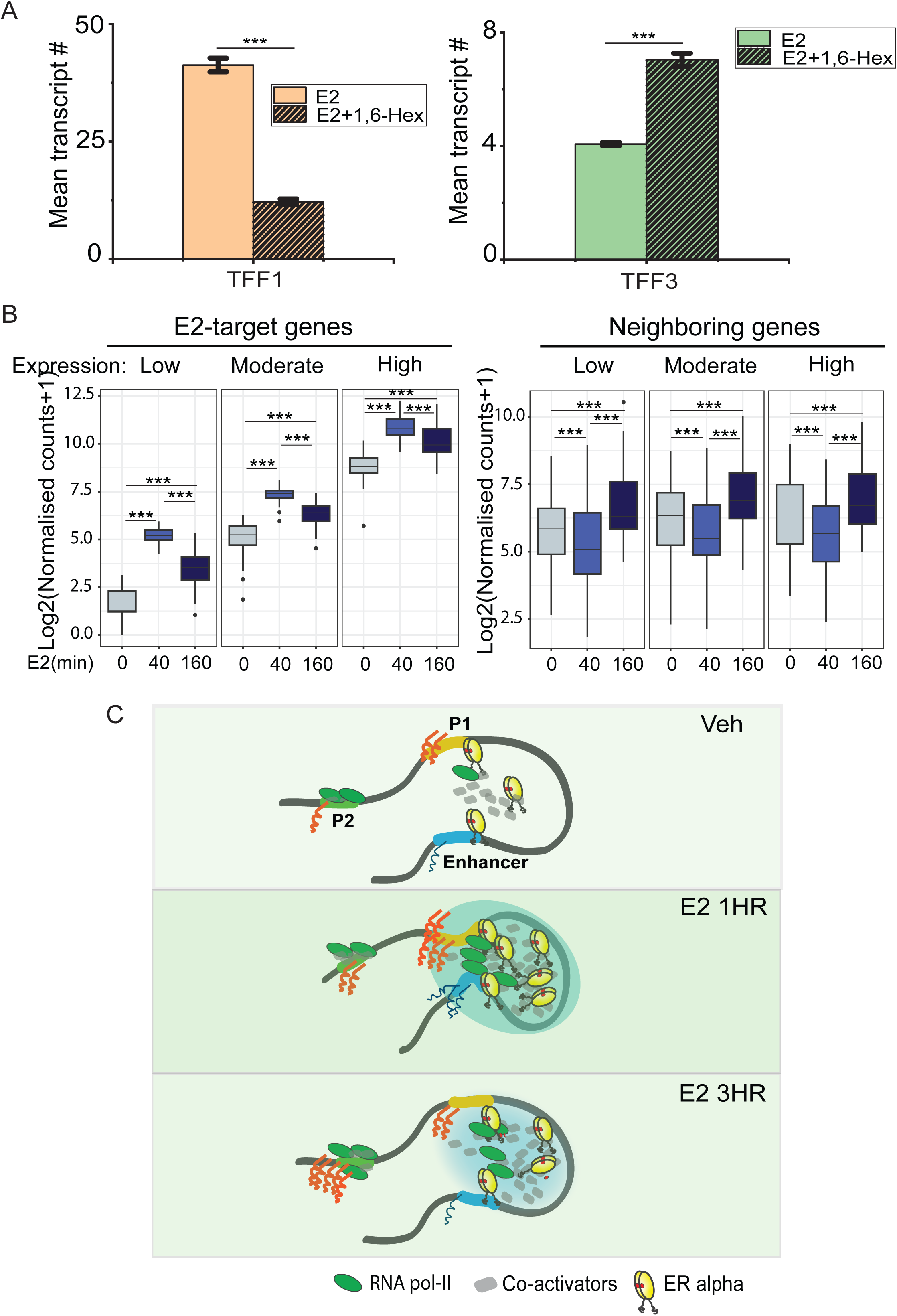
1,6 HD exposure down-regulates TFF1 but supports TFF3 expression. A. Mean transcript counts for InTFF1 and InTFF3 in control, and 3% 1,6-Hexanediol treated cells post 30 minutes of E2-induction. The mean of means are shown, and the error bars denote SEM from three repeats (n>880, N=3). p-values were calculated by Student’s two-tailed unpaired t-test, and the significance is represented as: *** denotes p < 0.001,** denotes p < 0.01,* denotes p < 0.05, ns denotes p > 0.05. B. Boxplots showing DESeq2 normalized counts for low expressing, moderately expressing, and highly-expressing genes in the vehicle, E2-40m, and E2-160m respectively (left). Boxplots showing DESeq2 normalized counts for genes near low expressing, moderately expressing, and highly expressing genes in the vehicle, E2-40m, and E2-160m, respectively (right). The p-values in the boxplots were calculated using the Wilcoxon rank-sum test. The boxplots depict the minimum, first quartile, median, third quartile, and maximum values, along with outliers. C. Model depicting the signaling under uninduced, E2-1hr, and E2-3hr conditions – First, activation of target gene loci (like *TFF1*) occurs by ligand-dependent induction. During the active phase (1 hr), liganded ERα binds on enhancer and promoter. Together these elements interact in 3D, manifesting as ERα punctae, which results in robust expression of target genes, and the sequestration of transcriptional machinery. At 3 hrs, as nuclear ERα decreases, the transcriptional machinery becomes available to other nearby promoters leading to increase in gene transcription at those loci.

## Discussion

smFISH allowed us to simultaneously capture allele level transcription of two genes, *TFF1* and *TFF3,* located in the same TAD, at single cell level during the peak and fall of signalling. We were able to capture the anti-correlated expression of these two genes, revealing intricate regulatory feedback between acutely activated enhancer-dependent gene, *TFF1,* which caused the dysregulation of the non-enhancer targeted gene, *TFF3,* within the same TAD (Figure 5B).

Our data suggests that while potential condensate formation on enhancer allows robust activation of the target gene, however, it negatively impacts the expression of other neighbouring genes possibly due to the sequestration of transcriptional machinery from these genes. When the condensates dissolve, the locally enriched transcriptional machinery is available to the other loci, allowing for increased transcription of these genes that are not the direct target of enhancers (Figure 5C). Forced dissolution of condensates by 1,6 HD allows the expression of non-enhancer target genes in the cells, although at low levels. The data also suggests that, globally, the genes next to ligand-induced genes are upregulated only at the fall of signalling as an indirect consequence of excess polymerase availability in the neighbourhood.

### Non-enhancer target genes are also regulated indirectly in the same TAD

ERα binding strength increases at the *TFF1* enhancer at 1h of ligand stimulation (Saravanan et al., 2020). Such estrogen-induced binding of ERα leads to robust ligand-induced activation of *TFF1* gene. While *TFF3* promoter remains unbound by ERα (Figure 3A) and does not interact with the *TFF1* enhancer throughout the course of signalling. These data suggest that TFF3 expression is both ERα and enhancer independent. Despite non-dependence, TFF3 expression was mutually exclusive to TFF1 expression and was negatively impacted in the absence of the enhancer. The latter could be due to the enhancer-mediated repositioning of the entire TAD that benefitted the expression of TFF3 and not due to the direct interaction between the enhancer and the *TFF3* gene.

The mean expression of TFF1 was manyfold higher than TFF3, thus explaining its dependence on an active upstream enhancer. The enhancer of *TFF1* interacts with the gene at the peak of signalling and shows some loss of interaction at 3h when signalling recedes. As stated above, the enhancer does not interact with *TFF3,* however, the enhancer forms a loop with *TFF3* upon deletion of the *TFF1* promoter (Oh et al., 2021). The ERα condensates are formed at the *TFF1* locus upon signalling (Saravanan et al., 2020), which is reduced upon enhancer deletion, suggesting its causal role in condensate formation. Corroborating with this, the enhancer of *TFF1* engages in the recruitment of megadalton complex (Liu et al., 2014) in MCF-7 cells, which could facilitate phase-separation (Nair et al., 2019, Saravanan et al., 2020). The cells that expressed TFF1 did not express TFF3 as efficiently, suggesting Pol2 was potentially sequestered from the *TFF3* promoter. However, at 3h when liganded ERα degrades leading to dissolution of potential condensates on enhancer, TFF3 expression increased, suggesting a local redistribution of active transcription machinery to other genes within the same chromatin domain. Even at 1h, when condensates were perturbed by 1,6 HD, the expression of TFF3 increased indicating the access to Pol2 pool by *TFF3* promoter. The data explains the negative correlation between TFF1 and TFF3 expression at the peak of signalling and robust activation of TFF3 at 3h when TFF1 expression decreased at single cell level. The presence of significant numbers of TFF1 nascent RNA in the nucleus in our data could also be due to a time lapse between transcription and splicing, as seen for the genes that are early responders during signaling (Zambrano et al., 2020).

### High and low levels of ERα were detrimental to enhancer mediated activation of *TFF1*

We observe that TFF1 expression is the highest in cells with medium levels of ERα, while it shows low expression in cells with low or high levels of ERα (Figure 4D, F). This points to an inverted U-shaped response of TFF1 to ERα, which is a nonmonotonic response to estrogen, as has been shown (Lebedeva et al., 2012; Vandenberg et al., 2012). A nonmonotonic response can be described as an inverted U-shaped curve with maximal response occurring in the middle and reducing substantially at very low or very high doses (Vandenberg et al., 2012). Further, we observe that for TFF3, the maximal expression occurs at low levels of ERα (Figure 4E, F). This could be indicative of a response such that a low concentration is enough to meet the threshold for expression beyond which all other higher doses attenuate the expression (Lebedeva et al., 2012; Vandenberg et al., 2012). This could also suggest that ERα levels have a varied effect on different genes and their expression depending on the extent of reliance of the gene on ERα for its expression. This phenomenon, called the prozone effect, is characterized by the inhibition of multimeric complexes when one of the constituents of the complex are present at very high concentrations as has been shown for TFF1 (Lebedeva et al., 2012). In the context of condensates on *TFF1* gene, we speculate that very low levels of TF may not reach or exceed the critical concentration required for LLPS (for *TFF1*), but this poor transcriptional state of *TFF1* could be favourable for *TFF3*. Alternatively, a very high concentration of the TF could result in the formation of highly dense homogenous condensates, effectively reducing transcription in a situation akin to the prozone effect (Chong et al., 2022; Lebedeva et al., 2012; Ryu et al., 2024).

Interestingly, the levels of ERα do not determine the expression of other E2 target genes like *GREB1* and *MYC* (Stossi et al., 2020); however, we observed that overexpression of ERα causes downregulation of TFF1 (Figure 4- figure supplement 1A) and this is in agreement with what has been observed for the expression status of *TFF1* in cell line with high ERα compared to cell lines with low ERα (Lebedeva et al., 2012).

ER is known to be expressed in nearly 70% of breast cancers, and many of these are treated using estrogen receptor modulators and degraders (Arnesen et al., 2021). Therefore, understanding the effect of ERα on the expression of estrogen-responsive genes depending on whether they are target genes or non-target genes, is of therapeutic importance as several non-target genes have been shown to be differentially regulated in ER-mutant breast cancer cells (Arnesen et al., 2021).

Together, our study underscores the indirect effects of ligand-induced chronic gene transcription that is dependent on enhancer activation and possible TF-condensate within a TAD.

## Materials and Methods

### Cell culture

MCF-7 cells (original line procured from ATCC; WT scramble control and Δ*TFF1*e were generated in Saravanan et al., 2020) were cultured in non-stripping media consisting of DMEM (Gibco, 12100-046) supplemented with 10% FBS (Gibco, 16000-044) and 1% Penicillin-Streptomycin-Glutamine (Gibco, 10378-016) in a 5% CO2 humidified incubator at 37 °C (unstripped MCF-7). Cells were passaged and seeded into glass bottom dishes in non-stripping media for 24h and allowed to reach 60–80% confluency. These cells were then hormone-stripped for three days in stripping media containing phenol red-free DMEM (Gibco, 21063-029) supplemented with 5% charcoal–stripped FBS (Gibco,12676029) and maintained in the humidified incubator at 37 °C (stripped MCF-7).

In order to induce the estrogen transcriptional responses, on the third day cells were treated with β-estradiol (E2758, Sigma-Aldrich) at 100nM concentration for various periods as mentioned in the respective Figures. For untreated control, cells were either treated with equal microliters of ethanol on the third day or with ERα inhibitor ICI182780 (1047, Tocris Biosciences) at 100nM concentration for 24 h after two days of stripping.

For perturbation of transcriptional condensates in E2-induced condition, cells were incubated with media containing 3% 1,6-Hexanediol for 5 minutes (after 30 minutes of E2 treatment) followed by removal of 1,6-Hexanediol and recovery for 25 mins in E2 containing media.

For experiments with overexpression of EGFP- ERα or EGFP, cells were transfected 24 hours before E2 induction using X-tremeGENE™ HP DNA Transfection Reagent (XTGHP-RO) in stripping media.

### Probe for smFISH

Probes were designed, custom-made, and tagged with indicated fluorophores (Quasar 570 or Cal Fluor 610) by Biosearch Technologies (https://www.biosearchtech.com/products/rna-fish). Probes were made to target the intronic or exonic regions of the *TFF1* and *TFF3* genes. Probe sequences can be found in supplementary file 1a-d in supplementary materials.

### Chromatin retention assay and smFISH

For smFISH experiments without chromatin retention assay cells, the cells were treated with estradiol or vehicular control for indicated times. Following this, the cells were washed with nuclease-free 1X PBS (Ambion, AM9624) twice. This was followed by fixation using 4% paraformaldehyde (PFA, Sigma, P6148) in 1X PBS for 10 minutes at room temperature. The fixative was removed and cells were washed twice with 1X PBS. The cells were permeabilized using 70% ethanol at 4°C overnight. On the next day, the permeabilizing agent was removed and the cells were washed twice with 1X PBS. The cells were washed once with a wash buffer (10% Formamide (Ambion, 9342) and 2X SSC (Ambion, AM9763) in nuclease-free water) for 5 minutes at room temperature. The wash buffer was aspirated and the cells were incubated with a hybridization mix (100 μl of mix contains 10% formamide, 89 μl hybridization buffer, and 1 μl each of indicated probes) overnight at 37°C in a humid chamber. The next day, the solution was removed and cells were washed with the wash buffer at 37°C for 30 minutes. To stain the nuclei, DAPI (Invitrogen, D1306; 2μg/ml) in wash buffer was added to the cells and incubated for 10 minutes at 37°C. The cells were then washed with 2X SSC for 5 minutes at 4°C. The solution was removed and the cells were covered with a few drops of the mountant Vectashield (Vector Labs). The plates were imaged after at least 1h.

For smFISH experiments with chromatin retention assay cells were treated with estradiol or vehicular control for indicated times. Following this, the cells were treated with CSK buffer (consisting of 10 mM PIPES buffer, 100 mM NaCl, 3 mM MgCl2, 300 mM sucrose, and 0.7% Triton-X 100) for 15 minutes at room temperature. After this, the cells were washed with nuclease-free 1X PBS twice. This was followed by fixation using 4% paraformaldehyde in 1X PBS for 10 minutes at room temperature. The fixative was removed and cells were washed twice with 1X PBS. The cells were permeabilized using 0.3% Triton-X 100 (Sigma, T8787) in 1X PBS for 10 minutes at room temperature. The permeabilizing agent was aspirated and the cells were washed twice with 1X PBS. This was followed by a washing step using the wash buffer for 5 minutes at room temperature. Following this, the cells were incubated with the hybridization mix to which the antibody of interest was also added at appropriate dilution. The next day, the hybridization mix containing probes and antibodies was removed. Cells were washed twice with 1X PBS. Then the cells were incubated with 1X PBS containing a secondary antibody against the antibody of interest at room temperature for 2h. This was followed by incubation with the wash buffer at 37°C for 30 minutes. To stain the nuclei, DAPI in wash buffer was added to the cells and incubated for 10 minutes at 37°C. The cells were then washed with 2X SSC for 5 minutes at 4°C. The solution was removed and the cells were covered with a few drops of the mountant Vectashield. The plates were imaged after at least 1h.

### Antibody staining/ immunofluorescence

The primary antibody against ERα (Santa Cruz, sc8002(F10)) was added to the hybridization mix at the dilution of 1:400 and incubated overnight at 37°C. The next day, an anti-mouse Alexa Fluor™ 488 secondary antibody (Invitrogen, A11029) was added to 1X PBS at the dilution of 1:1000. Cells were incubated for 2h followed by the continuation of the smFISH protocol as indicated above.

### Image acquisition

The plates were imaged on an Olympus IX83 inverted widefield fluorescence microscope with a Retiga 6000 CCD monochrome camera (QImaging). The images were acquired using a 60X, 1.42 N.A. oil immersion objective or a 100X, 1.4 N.A. oil immersion objective. The z-step size was 0.3μm and 35 slices were acquired. The resolution at which the images were acquired is 2752 x 2208. Narrow band-pass filters were used to distinguish the signal from Quasar 570 and Cal Fluor 610 labeled probes (ChromaTechnology- 49309 and 49310).

### Image analysis and representation

The images of mRNA channels were subjected to rolling ball background subtraction across the entire Z-stack using Fiji. For representative images, the stacks were Z-projected in Fiji. Transcripts were counted using either Rajlabimagetools (courtesy of Arjun Raj lab) or an in-house MATLAB (Mathworks) script based on earlier work (Femino et al., 1998; Raj et al., 2008). Briefly, the code uses a nuclear mask to count the intron-containing RNA present in the nucleus (InTFF), and another mask for the whole cell to count the mature mRNA, detected by exonic probes (ExTFF), that are present in the nucleus as well as the cytoplasm. The mean intensity for the ERα immunofluorescence channel was also quantified using the MATLAB script on a cell-by-cell basis using the nuclear mask.

### Graphing and Statistics

The graphs were plotted using Python 3, MATLAB, and Origin Pro and edited using Adobe Photoshop. To perform a student’s t-test for data combined from three repeats, GraphPad (https://www.graphpad.com/quickcalcs/ttest1/) was used. Non-parametric tests were performed in case of single cell data. For this Mann-Whitney test was used. Significance is represented as : *** denotes p < 0.001,** denotes p < 0.01,* denotes p < 0.05, ns denotes p > 0.05. The specific test used has been mentioned in the relevant figure legends.

### Circular Chromatin Conformation Capture-seq

4C was performed as per the protocol described in van de Werken et al., 2012 with minor variations. MCF7 cells were fixed with fresh formaldehyde (1.5%) and quenched with glycine (125mM) followed by washes with ice-cold 1XPBS (2X) and scraped, pelleted, and stored at - 80^0^C. Lysis buffer [Tris-Cl pH 8.0 (10mM), NaCl (10mM), NP-40 (0.2%), PIC (1X)] was added to the pellets and homogenized by Dounce homogenizer (20 stroked with pestle A followed by pestle B). The 3C digestion was performed with DpnII (200 units, NEB) and ligation was performed by T4 DNA ligase and ligation mix [Triton X-100 (1%), 1x Ligation buffer (10X Ligation buffer- Tris-Cl pH 7.5 (500mM), MgCl2 (100mM), DTT (100mM), BSA (0.105mg/ml), ATP (1.05mM)]. The ligated samples were purified by PCI and subjected to ethanol precipitation. The pellet was eluted in 1X TE (pH 8.0) to obtain the 3C library. The 4C digestion was performed by NlaIII (50 units, NEB), and the samples were ligated, purified, and precipitated similar to the 3C library to obtain the 4C library. The 4C library was subjected to RNaseA treatment and purified with the QIAquick PCR purification kit. The concentration of the library was then measured by Nanodrop and subjected to PCRs using the oligos for the enhancer viewpoint. The samples were next PCR purified using the same kit and subjected to next-generation sequencing with Illumina HiSeq2500/ NOVA seq. The 4C oligos are listed in supplementary file 1e.

### 4C Data Analysis

The sequenced reads in fastq file were demultiplexed by matching the appropriate primer sequences for each condition without allowing for any mismatches. Demultiplexed reads were processed using 4cseqpipe software. Restriction site tracks were created for the hg38 human genome by mentioning the restriction sites of the first cutter as GATC and that of the second cutter as CATG. The phred scores of demultiplexed reads were changed to phred64 format and the fastq files were converted into raw format. Further, valid 4C reads were mapped to the generated restriction site tracks. Unique fragment ends/non-unique fragment ends were used. The mapped reads were normalized and near cis domainograms at a maximum height of 0.1 were created by using the truncated log mean statistic with a trend resolution of 1kb for the genomic region chr11:42300000-42400000. Data given in (Fig 3A_source data 1 file).

### RNA Isolation, cDNA synthesis and PCR

Cells were lysed in 1 ml of Trizol (Thermo Fisher Inc.). 200 ul chloroform was added to the sample, briefly vortexed and centrifuged at 12K rpm for 12 mins. The aqueous phase was carefully collected and transferred to the fresh tube. 1 volume of isopropanol was added to the sample and incubated at room temperature for 10 mins to precipitate the RNA. The samples were centrifuged at 12K rpm for 12 mins, supernatant was discarded without disturbing the pellet. The pellet obtained was washed with 75% ethanol. The pellet was air dried and dissolved in RNase free water. RNA obtained was treated with ezDNase (Invitrogen) to remove the traces of contaminating DNA. 1μg of RNA was used for each cDNA synthesis reaction by Superscript IV (Invitrogen) and random hexamers as per manufacturer’s recommendation. The CFX96 touch (Biorad) real time PCR was used for qRT-PCRs. The fold changes were calculated by the ΔΔCt method and individual expression data was normalized to GAPDH mRNA. The *p*-values were calculated by Student’s unpaired two-tailed *t*-test from independent four biological replicates. qRT-PCR primers are listed in Supplementary file 1f.

### GRO-Seq analysis

Fastq files from GEO accession number GSE43836 were downloaded from European Nucleotide Archive. Reads with base quality <20 in a sliding window of 4 bases and with a length of <36 were removed using Trimmomatic 0.39. Trimmed reads were aligned using bowtie2 2.5.1 with default parameters. Duplicate reads from the alignment files were removed using samtools 1.16.1. De-duplicated aligned reads were assigned in a strand-specific manner to transcript feature of the hg38.ncbiRefSeq.gtf.gz file by allowing multi-mapping reads and considering the largest overlap in case of overlapping features using featureCounts v2.0.3. CPM normalised strand-specific signal files with a bin size of 1 bp were generated using bamCoverage 3.5.1. Differential gene expression analysis was performed with default parameters using Deseq2 1.36.0. Upregulated genes were defined as those with adjusted p-value <0.05 and log2(FC) >1 in their respective conditions. Top, Middle and bottom 10% of upregulated genes based on their base mean value in E2-40m vs VEH condition were subsetted as highly expressing, moderately expressing and low expressing upregulated genes. The closest upregulated genes in E2-160m vs E2-40m near highly expressing, moderately expressing and low expressing upregulated genes in E2-40m vs VEH condition were identified using bedtools closest v2.30.0. Only genes within a genomic distance of 1Mb were considered. Log2 transformed DEseq2 normalised counts with the addition of the arbitrary value 1 was used to compare the gene expression trends across time points and categories of upregulated genes. All plots were generated using R 4.2.2.

## Funding and Acknowledgements

This project was supported by intramural funds at TIFR Hyderabad from the Department of Atomic Energy, Government of India (Project Identification No. RTI 4007). We further acknowledge support of the Department of Atomic Energy, Government of India, under project no. 12-R&D-TFR-5.04-0800 and intramural funds from National Centre for Biological Sciences–Tata Institute of Fundamental Research (NCBS-TIFR). D.N. is an EMBO Global Investigator. We also acknowledge the funding support from Wellcome-IA(IA/1/14/2/501539 and IA/S/23/1/506749). DB is supported by TIFR-Hyderabad PhD program. ZI acknowledges funding support from ICMR, India. SN acknowledges funding support from DBT-JRF, India.

## Material and Data Availability

All source data associated with the manuscript are provided as supplements. Probe and primer sequences are provided as supplementary tables. These are listed below.

---

**Figure 1—figure supplement 1. Experimental design**

A. Schematic depicting the experimental design. Cells were cultured in complete media for 24h followed by stripping for 3 days. Finally, cells were induced for different durations with E2/Vehicle followed by different assays like smFISH, 4C-seq, smFISH-IF, etc.

B. Zoomed in region around TFF3 gene, UCSC genome browser snapshots showing the binding of ERα and H3k4me3 signal. First, second and third ERα ChIP-seq tracks are in vehicle treated, E2 treated for 1h and E2 treated for 3h in WT cells respectively

C. UCSC genome browser snapshots showing Gro-seq signal for *TFF3* gene. First, second and third Gro-seq tracks are from vehicle-treated, E2-40m and E2-160m in WT cells, respectively.

**Figure 2—figure supplement 1. Dynamic induction and RNA localization of *TFF1* and *TFF3* transcription across cell populations using smRNA FISH**

A. Bar graph depicting the percentage of cells with 1,2,3,4, or greater than 4 sites of transcription for TFF1 (left) is shown. The graph shows the mean of means from different repeats of the experiment, and error bars denote SEM (n>200, N=3). Only the cells with at least one allele firing were counted and cells with no alleles were not included in this. The graph on right shows the number of cells with zero or non-zero number of alleles firing.

B. Bar graph depicting the percentage of cells with 1,2,3,4 or greater than 4 sites of transcription for TFF3 (left) is shown. The graph shows the mean of means from different repeats of the experiment, and error bars denote SEM (n>200, N=3). Only the cells with at least one allele firing were counted and cells with no alleles were not included in this. The graph in the middle shows the number of cells with 2,3,4 or greater than 4 sites of transcription for TFF3.The graph on the right shows the number of cells with zero or non-zero number of alleles firing.

C. Images from single molecule RNA FISH experiment showing transcripts for InTFF1 in cells induced for 1 hour with E2. The image shows that when a single allele of TFF1 is firing, the transcripts show a more spatially restricted localisation. The scale bar is 5 microns.

D. Images from single molecule RNA FISH experiment showing transcripts for InTFF1 in uninduced cells. The image shows that when a single allele of TFF1 is firing and transcription is low, the transcripts show a more spatially restricted localisation. The scale bar is 5 microns.

E. 60X Representative images from a single molecule RNA FISH experiment showing transcripts for InTFF1 and ExTFF1 (top) and InTFF3 and ExTFF3 (bottom). The image shows that there is no intronic signal in the cytoplasm, while exonic signals can be found both in the nucleus and the cytoplasm. The scale bar is 5 microns.

F. 60X Representative images from single molecule RNA FISH experiment showing transcripts for InTFF1 and ExTFF1. The image shows that all intronic signals are colocalized with exonic signals, but all exonic signals are expectedly not colocalized with intronic signals, representing more mature mRNA. The scale bar is 5 microns.

**Figure 2—figure supplement 2. *TFF1* and *TFF3* expressions show opposite trends during the E2-signaling time-course**

A. Violin plot showing the ratio of total intronic to absolute exonic *TFF1* counts are depicted. Absolute exonic counts are calculated by subtracting total intronic transcripts from total exonic transcripts. The graph shows the distribution of ratios combined from three different repeats. Error bars denote SEM. p-values were calculated by the Mann-Whitney test, and the significance is represented as: *** denotes p < 0.001, ** denotes p < 0.01, * denotes p < 0.05, and ns denotes p > 0.05.

B. Violin plot showing the ratio of total intronic to absolute exonic *TFF3* counts are depicted. Absolute exonic counts are calculated by subtracting total intronic transcripts from total exonic transcripts. The graph shows the distribution of ratios combined from three different repeats. Error bars denote SEM. p-values were calculated by the Mann-Whitney test, and the significance is represented as: *** denotes p < 0.001, ** denotes p < 0.01, * denotes p < 0.05, and ns denotes p > 0.05.

**Figure 3—figure supplement 1. *TFF1* enhancer does not change target promoters during signalling time-course**

4C-seq plot at *TFF1* enhancer viewpoint, the interaction with the promoter is highlighted in yellow. The plot is overlaid with H3K27ac, ERα ChIP signal, and gene annotations. There is no substantial interaction between the enhancer and *TFF3* locus at any time point, while the interaction between the enhancer and *TFF1* locus increases at 1hr and decreases at 3hr.

**Figure 4—figure supplement 1. Levels of ERα in the nucleus dictate the extent of *TFF1* and *TFF3* inductions**

A. Representative images from smFISH experiments showing InTFF1 and GAPDH (top panel) and InTFF1 and InTFF3 (bottom panel) in cells overexpressing ERα-GFP. Yellow circles show cells that are GFP positive, and the transcripts associated with them. Note the visibly fewer TFF1, and TFF3 transcripts in the ERα-GFP positive cells, while GAPDH transcripts remain indistinguishable. This shows that ERα-GFP overexpression specifically affects the transcription of E2-regulated genes like TFF1 and TFF3 and not a housekeeping gene like GAPDH. This is further quantified in B and C.

B. Scatter plots showing the distribution of ERα-GFP intensities with GAPDH or InTFF1 transcript counts on a cell-by-cell basis from the experiment in the first row of A. Cells with high ERα-GFP expression can have high GAPDH expression, but such cells necessarily have low TFF1 counts.

C. Scatter plots showing the distribution of ERα GFP intensities with InTFF1 or InTFF3 counts on a cell-by-cell basis from the experiment in the second row of A. As in B, high ERα-GFP expression leads to low expression of E2-regulated genes like *TFF1* or *TFF3*.

D. Representative images showing smFISH for InTFF1 and InTFF3 in cells overexpressing EGFP-C1. Yellow circles show cells that are GFP positive, and the transcripts associated with them. Just expressing EGFP in the same backbone as the ERα-GFP plasmid, does not downregulate *TFF1*, and *TFF3* expression; indicating that the downregulation effect is specific to ERα overexpression and not a generic effect of cell transfection. This is quantified in E.

E. Scatter plots showing the distribution of EGFP-C1 intensities with InTFF1 or InTFF3 transcript counts on a cell-by-cell basis. There is no obvious correlation between the levels of EGFP-C1 and TFF1 or TFF3, and a cell with high EGFP levels may well have large TFF1 or TFF3 transcript counts.

**Figure 5—figure supplement 1. LLPS perturbation down-regulates *TFF1* but supports *TFF3* expression**

Mean transcript counts for InTFF1 and InTFF3 in control, and 3% 1,6-Hexanediol treated cells in the absence of E2-induction show no difference in TFF1 counts on 1,6-Hexanediol treatment. The bar denotes the mean of means from two repeats, and the error bars denote SEM. p-values were calculated by Student’s two-tailed unpaired t-test, and the significance is represented as: *** denotes p < 0.001, ** denotes p < 0.01, * denotes p < 0.05, ns denotes p > 0.05.

## Supplementary data files

Supplementary File 1a- InTFF3 probe sequence Supplementary File 1b- ExTFF3 probe sequence Supplementary File 1c- InTFF1 probe sequence Supplementary File 1d- ExTFF1 probe sequence Supplementary File 1e- 4C Seq oligo sequences Supplementary File 1f- qPCR oligo sequences

## Source data files

Figure 1- source data 1

Figure 2- source data 1

Figure 2-Figure Supplement 1- source data 1

Figure 2-Figure Supplement 2- source data 1

Figure 3- source data 1

Figure 3A-source data 1

Figure 3-Figure Supplement 1- source data 1

Figure 4- source data 1

Figure 4-Figure Supplement 1- source data 1

Figure 5- source data 1

Figure 5-Figure Supplement 1- source data 1

## References

Ahmed, J., Meszaros, A., Lazar, T., & Tompa, P. (2021). DNA-binding domain as the minimal region driving RNA-dependent liquid-liquid phase separation of androgen receptor. Protein Science, 30(7), 1380–1392. 10.1002/pro.4100

Arnesen, S., Blanchard, Z., Williams, M. M., Berrett, K. C., Li, Z., Oesterreich, S., Richer, J. K., & Gertz, J. (2021). Estrogen Receptor Alpha Mutations in Breast Cancer Cells Cause Gene Expression Changes through Constant Activity and Secondary Effects. Cancer Research, 81(3), 539–551. 10.1158/0008-5472.CAN-20-1171

Boehning, M., Dugast-Darzacq, C., Rankovic, M., Hansen, A. S., Yu, T., Marie-Nelly, H., McSwiggen, D. T., Kokic, G., Dailey, G. M., Cramer, P., Darzacq, X., & Zweckstetter, M. (2018). RNA polymerase II clustering through carboxy-terminal domain phase separation. Nature Structural & Molecular Biology, 25(9), 833–840. 10.1038/s41594-018-0112-y

Boija, A., Klein, I. A., Sabari, B. R., Dall’Agnese, A., Coffey, E. L., Zamudio, A. V., Li, C. H., Shrinivas, K., Manteiga, J. C., Hannett, N. M., Abraham, B. J., Afeyan, L. K., Guo, Y. E., Rimel, J. K., Fant, C. B., Schuijers, J., Lee, T. I., Taatjes, D. J., & Young, R. A. (2018). Transcription Factors Activate Genes through the Phase-Separation Capacity of Their Activation Domains. Cell, 175(7), 1842–1855.e16. 10.1016/j.cell.2018.10.042

Bojcsuk, D., Nagy, G., & Balint, B. L. (2017). Inducible super-enhancers are organized based on canonical signal-specific transcription factor binding elements. Nucleic Acids Research, 45(7), 3693–3706. 10.1093/nar/gkw1283

Brown, A. M., Jeltsch, J. M., Roberts, M., & Chambon, P. (1984). Activation of pS2 gene transcription is a primary response to estrogen in the human breast cancer cell line MCF-7. Proceedings of the National Academy of Sciences, 81(20), 6344–6348. 10.1073/pnas.81.20.6344

Buecker, C., Srinivasan, R., Wu, Z., Calo, E., Acampora, D., Faial, T., Simeone, A., Tan, M., Swigut, T., & Wysocka, J. (2014). Reorganization of enhancer patterns in transition from naive to primed pluripotency. Cell Stem Cell, 14(6), 838–853. 10.1016/j.stem.2014.04.003

Bulger, M., & Groudine, M. (1999). Looping versus linking: Toward a model for long-distance gene activation. Genes & Development, 13(19), 2465–2477.

Cai, D., Feliciano, D., Dong, P., Flores, E., Gruebele, M., Porat-Shliom, N., Sukenik, S., Liu, Z., & Lippincott-Schwartz, J. (2019). Phase separation of YAP reorganizes genome topology for long-term YAP target gene expression. Nature Cell Biology, 21(12), 1578–1589. 10.1038/s41556-019-0433-z

Chalbos, D., Philips, A., Galtier, F., & Rochefort, H. (1993). Synthetic antiestrogens modulate induction of pS2 and cathepsin-D messenger ribonucleic acid by growth factors and adenosine 3’,5’-monophosphate in MCF7 cells. Endocrinology, 133(2), 571–576. 10.1210/en.133.2.571

Chen, H., Levo, M., Barinov, L., Fujioka, M., Jaynes, J. B., & Gregor, T. (2018). Dynamic interplay between enhancer-promoter topology and gene activity. Nature Genetics, 50(9), 1296–1303. 10.1038/s41588-018-0175-z

Chen, L., Zhang, Z., Han, Q., Maity, B. K., Rodrigues, L., Zboril, E., Adhikari, R., Ko, S.-H., Li, X., Yoshida, S. R., Xue, P., Smith, E., Xu, K., Wang, Q., Huang, T. H.-M., Chong, S., & Liu, Z. (2023). Hormone-induced enhancer assembly requires an optimal level of hormone receptor multivalent interactions. Molecular Cell, 83(19), 3438–3456.e12. 10.1016/j.molcel.2023.08.027

Chepelev, I., Wei, G., Wangsa, D., Tang, Q., & Zhao, K. (2012). Characterization of genome-wide enhancer-promoter interactions reveals co-expression of interacting genes and modes of higher order chromatin organization. Cell Research, 22(3), 490–503. 10.1038/cr.2012.15

Chinery, R., Williamson, J., & Poulsom, R. (1996). The Gene Encoding Human Intestinal Trefoil Factor (*TFF3*) Is Located on Chromosome 21q22.3 Clustered with Other Members of the Trefoil Peptide Family. Genomics, 32(2), 281–284. 10.1006/geno.1996.0117

Cho, W.-K., Spille, J.-H., Hecht, M., Lee, C., Li, C., Grube, V., & Cisse, I. I. (2018). Mediator and RNA polymerase II clusters associate in transcription-dependent condensates. Science, 361(6400), 412–415. 10.1126/science.aar4199

Chong, S., Dugast-Darzacq, C., Liu, Z., Dong, P., Dailey, G. M., Cattoglio, C., Heckert, A., Banala, S., Lavis, L., Darzacq, X., & Tjian, R. (2018). Imaging dynamic and selective low-complexity domain interactions that control gene transcription. Science, 361(6400), eaar2555. 10.1126/science.aar2555

Chong, S., Graham, T. G. W., Dugast-Darzacq, C., Dailey, G. M., Darzacq, X., & Tjian, R. (2022). Tuning levels of low-complexity domain interactions to modulate endogenous oncogenic transcription. Molecular Cell, 82(11), 2084–2097.e5. 10.1016/j.molcel.2022.04.007

Clark, M. B., Mercer, T. R., Bussotti, G., Leonardi, T., Haynes, K. R., Crawford, J., Brunck, M. E., Cao, K.-A. L., Thomas, G. P., Chen, W. Y., Taft, R. J., Nielsen, L. K., Enright, A. J., Mattick, J. S., & Dinger, M. E. (2015). Quantitative gene profiling of long noncoding RNAs with targeted RNA sequencing. Nature Methods, 12(4), 339–342. 10.1038/nmeth.3321

Conesa, A., Madrigal, P., Tarazona, S., Gomez-Cabrero, D., Cervera, A., McPherson, A., Szcześniak, M. W., Gaffney, D. J., Elo, L. L., Zhang, X., & Mortazavi, A. (2016). A survey of best practices for RNA-seq data analysis. Genome Biology, 17(1), 13. 10.1186/s13059-016-0881-8

Coté, A., O’Farrell, A., Dardani, I., Dunagin, M., Coté, C., Wan, Y., Bayatpour, S., Drexler, H. L., Alexander, K. A., Chen, F., Wassie, A. T., Patel, R., Pham, K., Boyden, E. S., Berger, S., Phillips-Cremins, J., Churchman, L. S., & Raj, A. (2023). Post-transcriptional splicing can occur in a slow-moving zone around the gene. eLife, 12. 10.7554/eLife.91357.2

Coulon, A., Ferguson, M. L., de Turris, V., Palangat, M., Chow, C. C., & Larson, D. R. (2014). Kinetic competition during the transcription cycle results in stochastic RNA processing. eLife, 3, e03939. 10.7554/eLife.03939

Dixon, B. S., Sharma, R. V., & Dennis, M. J. (1996). The Bradykinin B2 Receptor Is a Delayed Early Response Gene for Platelet-derived Growth Factor in Arterial Smooth Muscle Cells *. Journal of Biological Chemistry, 271(23), 13324–13332. 10.1074/jbc.271.23.13324

Drexler, H. L., Choquet, K., & Churchman, L. S. (2020). Splicing Kinetics and Coordination Revealed by Direct Nascent RNA Sequencing through Nanopores. Molecular Cell, 77(5), 985–998.e8. 10.1016/j.molcel.2019.11.017

Femino, A. M., Fay, F. S., Fogarty, K., & Singer, R. H. (1998). Visualization of single RNA transcripts in situ. Science (New York, N.Y.), 280(5363), 585–590. 10.1126/science.280.5363.585

Fowler, T., Sen, R., & Roy, A. L. (2011). Regulation of Primary Response Genes. Molecular Cell, 44(3), 348–360. 10.1016/j.molcel.2011.09.014

Freter, R. R., Alberta, J. A., Hwang, G. Y., Wrentmore, A. L., & Stiles, C. D. (1996). Platelet-derived Growth Factor Induction of the Immediate-early gene MCP-1 Is Mediated by NF-κB and a 90-kDa Phosphoprotein Coactivator *. Journal of Biological Chemistry, 271(29), 17417–17424. 10.1074/jbc.271.29.17417

Friedman, M. J., Wagner, T., Lee, H., Rosenfeld, M. G., & Oh, S. (2024). Enhancer–promoter specificity in gene transcription: Molecular mechanisms and disease associations. Experimental & Molecular Medicine, 56(4), 772–787. 10.1038/s12276-024-01233-y

Fullwood, M. J., Liu, M. H., Pan, Y. F., Liu, J., Xu, H., Mohamed, Y. B., Orlov, Y. L., Velkov, S., Ho, A., Mei, P. H., Chew, E. G. Y., Huang, P. Y. H., Welboren, W.-J., Han, Y., Ooi, H. S., Ariyaratne, P. N., Vega, V. B., Luo, Y., Tan, P. Y., … Ruan, Y. (2009). An oestrogen-receptor-α-bound human chromatin interactome. Nature, 462(7269), Article 7269. 10.1038/nature08497

Furlong, E. E. M., & Levine, M. (2018). Developmental enhancers and chromosome topology. Science, 361(6409), 1341–1345. 10.1126/science.aau0320

Galouzis, C. C., & Furlong, E. E. M. (2022). Regulating specificity in enhancer-promoter communication. Current Opinion in Cell Biology, 75, 102065. 10.1016/j.ceb.2022.01.010

Gamliel, A., Meluzzi, D., Oh, S., Jiang, N., Destici, E., Rosenfeld, M. G., & Nair, S. J. (2022). Long-distance association of topological boundaries through nuclear condensates. Proceedings of the National Academy of Sciences of the United States of America, 119(32), e2206216119. 10.1073/pnas.2206216119

Hah, N., Danko, C. G., Core, L., Waterfall, J. J., Siepel, A., Lis, J. T., & Kraus, W. L. (2011). A rapid, extensive, and transient transcriptional response to estrogen signaling in breast cancer cells. Cell, 145(4), 622–634. 10.1016/j.cell.2011.03.042

Hah, N., Murakami, S., Nagari, A., Danko, C. G., & Kraus, W. L. (2013). Enhancer transcripts mark active estrogen receptor binding sites. Genome Research, 23(8), 1210–1223. 10.1101/gr.152306.112

Haimovich, G., & Gerst, J. E. (2018). Single-molecule Fluorescence in situ Hybridization (smFISH) for RNA Detection in Adherent Animal Cells. Bio-Protocol, 8(21), e3070. 10.21769/BioProtoc.3070

Herschman, H. R. (1991). Primary response genes induced by growth factors and tumor promoters. Annual Review of Biochemistry, 60(Volume 60, 1991), 281–319. 10.1146/annurev.bi.60.070191.001433

Jeltsch, J. M., Roberts, M., Schatz, C., Garnier, J. M., Brown, A. M., & Chambon, P. (1987). Structure of the human oestrogen-responsive gene pS2. Nucleic Acids Research, 15(4), 1401–1414.

Kagey, M. H., Newman, J. J., Bilodeau, S., Zhan, Y., Orlando, D. A., van Berkum, N. L., Ebmeier, C. C., Goossens, J., Rahl, P. B., Levine, S. S., Taatjes, D. J., Dekker, J., & Young, R. A. (2010). Mediator and cohesin connect gene expression and chromatin architecture. Nature, 467(7314), 430–435. 10.1038/nature09380

Khodor, Y. L., Menet, J. S., Tolan, M., & Rosbash, M. (2012). Cotranscriptional splicing efficiency differs dramatically between Drosophila and mouse. RNA, 18(12), 2174–2186. 10.1261/rna.034090.112

Kim, J., Petz, L. N., Ziegler, Y. S., Wood, J. R., Potthoff, S. J., & Nardulli, A. M. (2000). Regulation of the estrogen-responsive pS2 gene in MCF-7 human breast cancer cells. The Journal of Steroid Biochemistry and Molecular Biology, 74(4), 157–168. 10.1016/S0960-0760(00)00119-9

Koşar, Z., & Erbaş, A. (2022). Can the Concentration of a Transcription Factor Affect Gene Expression? Frontiers in Soft Matter, 2. https://www.frontiersin.org/articles/10.3389/frsfm.2022.914494

Kroschwald, S., Maharana, S., & Simon, A. (2017, May 22). Hexanediol: A chemical probe to investigate the material properties of membrane-less compartments. Matters. 10.19185/matters.201702000010

Kwon, S. (2013). Single-molecule fluorescence in situ hybridization: Quantitative imaging of single RNA molecules. BMB Reports, 46(2), 65–72. 10.5483/BMBRep.2013.46.2.016

Lebedeva, G., Yamaguchi, A., Langdon, S. P., Macleod, K., & Harrison, D. J. (2012). A model of estrogen-related gene expression reveals non-linear effects in transcriptional response to tamoxifen. BMC Systems Biology, 6(1), 138. 10.1186/1752-0509-6-138

Levenson, A. S., & Jordan, V. C. (1997). MCF-7: The First Hormone-responsive Breast Cancer Cell Line1. Cancer Research, 57(15), 3071–3078.

Levine, M., & Tjian, R. (2003). Transcription regulation and animal diversity. Nature, 424(6945), 147–151. 10.1038/nature01763

Li, W., Notani, D., Ma, Q., Tanasa, B., Nunez, E., Chen, A. Y., Merkurjev, D., Zhang, J., Ohgi, K., Song, X., Oh, S., Kim, H.-S., Glass, C. K., & Rosenfeld, M. G. (2013). Functional roles of enhancer RNAs for oestrogen-dependent transcriptional activation. Nature, 498(7455), 516–520. 10.1038/nature12210

Liu, Z., Merkurjev, D., Yang, F., Li, W., Oh, S., Friedman, M. J., Song, X., Zhang, F., Ma, Q., Ohgi, K. A., Krones, A., & Rosenfeld, M. G. (2014). Enhancer Activation Requires trans-Recruitment of a Mega Transcription Factor Complex. Cell, 159(2), 358–373. 10.1016/j.cell.2014.08.027

Mann, R., & Notani, D. (2023). Transcription factor condensates and signaling driven transcription. Nucleus, 14(1), 2205758. 10.1080/19491034.2023.2205758

Masiakowski, P., Breathnach, R., Bloch, J., Gannon, F., Krust, A., & Chambon, P. (1982). Cloning of cDNA sequences of hormone-regulated genes from the MCF-7 human breast cancer cell line. Nucleic Acids Research, 10(24), 7895–7903. 10.1093/nar/10.24.7895

Nair, S. J., Yang, L., Meluzzi, D., Oh, S., Yang, F., Friedman, M. J., Wang, S., Suter, T., Alshareedah, I., Gamliel, A., Ma, Q., Zhang, J., Hu, Y., Tan, Y., Ohgi, K. A., Jayani, R. S., Banerjee, P. R., Aggarwal, A. K., & Rosenfeld, M. G. (2019). Phase separation of ligand-activated enhancers licenses cooperative chromosomal enhancer assembly. Nature Structural & Molecular Biology, 26(3), Article 3. 10.1038/s41594-019-0190-5

Nunez, A. M., Berry, M., Imler, J. L., & Chambon, P. (1989). The 5’ flanking region of the pS2 gene contains a complex enhancer region responsive to oestrogens, epidermal growth factor, a tumour promoter (TPA), the c-Ha-ras oncoprotein and the c-jun protein. The EMBO Journal, 8(3), 823–829.

Oh, S., Shao, J., Mitra, J., Xiong, F., D’Antonio, M., Wang, R., Garcia-Bassets, I., Ma, Q., Zhu, X., Lee, J.-H., Nair, S. J., Yang, F., Ohgi, K., Frazer, K. A., Zhang, Z. D., Li, W., & Rosenfeld, M. G. (2021). Enhancer release and retargeting activates disease-susceptibility genes. Nature, 595(7869), Article 7869. 10.1038/s41586-021-03577-1

Panigrahi, A., & O’Malley, B. W. (2021). Mechanisms of enhancer action: The known and the unknown. Genome Biology, 22(1), Article 1. 10.1186/s13059-021-02322-1

Patange, S., Ball, D. A., Wan, Y., Karpova, T. S., Girvan, M., Levens, D., & Larson, D. R. (2022). MYC amplifies gene expression through global changes in transcription factor dynamics. Cell Reports, 38(4). 10.1016/j.celrep.2021.110292

Ptashne, M. (1986). Gene regulation by proteins acting nearby and at a distance. Nature, 322(6081), 697–701. 10.1038/322697a0

Quintin, J., Le Péron, C., Palierne, G., Bizot, M., Cunha, S., Sérandour, A. A., Avner, S., Henry, C., Percevault, F., Belaud-Rotureau, M.-A., Huet, S., Watrin, E., Eeckhoute, J., Legagneux, V., Salbert, G., & Métivier, R. (2014). Dynamic Estrogen Receptor Interactomes Control Estrogen-Responsive Trefoil Factor (TFF) Locus Cell-Specific Activities. Molecular and Cellular Biology, 34(13), 2418–2436. 10.1128/MCB.00918-13

Raj, A., Peskin, C. S., Tranchina, D., Vargas, D. Y., & Tyagi, S. (2006). Stochastic mRNA Synthesis in Mammalian Cells. PLOS Biology, 4(10), e309. 10.1371/journal.pbio.0040309

Raj, A., van den Bogaard, P., Rifkin, S. A., van Oudenaarden, A., & Tyagi, S. (2008). Imaging individual mRNA molecules using multiple singly labeled probes. Nature Methods, 5(10), Article 10. 10.1038/nmeth.1253

Rao, S.S.P. et al. (2014) ‘A 3D map of the human genome at kilobase resolution reveals principles of chromatin looping’, Cell, 159(7), pp. 1665–1680. doi:10.1016/j.cell.2014.11.021.

Richter, W. F., Nayak, S., Iwasa, J., & Taatjes, D. J. (2022). The Mediator complex as a master regulator of transcription by RNA polymerase II. Nature Reviews Molecular Cell Biology, 23(11), Article 11. 10.1038/s41580-022-00498-3

Rodriguez, J., Ren, G., Day, C. R., Zhao, K., Chow, C. C., & Larson, D. R. (2019). Intrinsic Dynamics of a Human Gene Reveal the Basis of Expression Heterogeneity. Cell, 176(1–2), 213–226.e18. 10.1016/j.cell.2018.11.026

Ryu, K., Park, G., & Cho, W.-K. (2024). Emerging insights into transcriptional condensates. Experimental & Molecular Medicine, 56(4), 820–826. 10.1038/s12276-024-01228-9

Sabari, B. R., Dall’Agnese, A., Boija, A., Klein, I. A., Coffey, E. L., Shrinivas, K., Abraham, B. J., Hannett, N. M., Zamudio, A. V., Manteiga, J. C., Li, C. H., Guo, Y. E., Day, D. S., Schuijers, J., Vasile, E., Malik, S., Hnisz, D., Lee, T. I., Cisse, I. I., … Young, R. A. (2018). Coactivator condensation at super-enhancers links phase separation and gene control. Science, 361(6400), eaar3958. 10.1126/science.aar3958

Sanyal, A., Lajoie, B. R., Jain, G., & Dekker, J. (2012). The long-range interaction landscape of gene promoters. Nature, 489(7414), 109–113. 10.1038/nature11279

Saravanan, B., Soota, D., Islam, Z., Majumdar, S., Mann, R., Meel, S., Farooq, U., Walavalkar, K., Gayen, S., Singh, A. K., Hannenhalli, S., & Notani, D. (2020). Ligand dependent gene regulation by transient ERα clustered enhancers. PLOS Genetics, 16(1), e1008516. 10.1371/journal.pgen.1008516

Sha, Y., Phan, J. H., & Wang, M. D. (2015). Effect of low-expression gene filtering on detection of differentially expressed genes in RNA-seq data. Conference Proceedings : … Annual International Conference of the IEEE Engineering in Medicine and Biology Society. IEEE Engineering in Medicine and Biology Society. Annual Conference, 2015, 6461. 10.1109/EMBC.2015.7319872

Shi, Y., Chen, J., Zeng, W.-J., Li, M., Zhao, W., Zhang, X.-D., & Yao, J. (2021). Formation of nuclear condensates by the Mediator complex subunit Med15 in mammalian cells. BMC Biology, 19(1), 245. 10.1186/s12915-021-01178-y

Shrinivas, K., Sabari, B. R., Coffey, E. L., Klein, I. A., Boija, A., Zamudio, A. V., Schuijers, J., Hannett, N. M., Sharp, P. A., Young, R. A., & Chakraborty, A. K. (2019). Enhancer Features that Drive Formation of Transcriptional Condensates. Molecular Cell, 75(3), 549–561.e7. 10.1016/j.molcel.2019.07.009

Skinner, S. O., Xu, H., Nagarkar-Jaiswal, S., Freire, P. R., Zwaka, T. P., & Golding, I. (2016). Single-cell analysis of transcription kinetics across the cell cycle. eLife, 5, e12175. 10.7554/eLife.12175

Stortz, M., Pecci, A., Presman, D. M., & Levi, V. (2020). Unraveling the molecular interactions involved in phase separation of glucocorticoid receptor. BMC Biology, 18(1), 59. 10.1186/s12915-020-00788-2

Stortz, M., Presman, D. M., & Levi, V. (2024). Transcriptional condensates: A blessing or a curse for gene regulation? Communications Biology, 7(1), 1–10. 10.1038/s42003-024-05892-5

Stossi, F., Dandekar, R. D., Mancini, M. G., Gu, G., Fuqua, S. A. W., Nardone, A., De Angelis, C., Fu, X., Schiff, R., Bedford, M. T., Xu, W., Johansson, H. E., Stephan, C. C., & Mancini, M. A. (2020). Estrogen-induced transcription at individual alleles is independent of receptor level and active conformation but can be modulated by coactivators activity. Nucleic Acids Research, 48(4), 1800. 10.1093/nar/gkz1172

Sun, J.-M., Spencer, V. A., Li, L., Yu Chen, H., Yu, J., & Davie, J. R. (2005). Estrogen regulation of trefoil factor 1 expression by estrogen receptor α and Sp proteins. Experimental Cell Research, 302(1), 96–107. 10.1016/j.yexcr.2004.08.015

Svensson, V., Natarajan, K. N., Ly, L.-H., Miragaia, R. J., Labalette, C., Macaulay, I. C., Cvejic, A., & Teichmann, S. A. (2017). Power analysis of single-cell RNA-sequencing experiments. Nature Methods, 14(4), 381–387. 10.1038/nmeth.4220

Tarazona, S., García-Alcalde, F., Dopazo, J., Ferrer, A., & Conesa, A. (2011). Differential expression in RNA-seq: A matter of depth. Genome Research, 21(12), 2213–2223. 10.1101/gr.124321.111

Uyehara, C. M., & Apostolou, E. (2023). 3D enhancer-promoter interactions and multi-connected hubs: Organizational principles and functional roles. Cell Reports, 42(4), 112068. 10.1016/j.celrep.2023.112068

van de Werken HJ, de Vree PJ, Splinter E, Holwerda SJ, Klous P, de Wit E, de Laat W. 4C technology: protocols and data analysis. Methods Enzymol. 2012;513:89–112. doi: 10.1016/B978-0-12-391938-0.00004-5. PMID: 22929766.

Vandenberg, L. N., Colborn, T., Hayes, T. B., Heindel, J. J., Jacobs, D. R., Jr., Lee, D.-H., Shioda, T., Soto, A. M., vom Saal, F. S., Welshons, W. V., Zoeller, R. T., & Myers, J. P. (2012). Hormones and Endocrine-Disrupting Chemicals: Low-Dose Effects and Nonmonotonic Dose Responses. Endocrine Reviews, 33(3), 378–455. 10.1210/er.2011-1050

Williams, M., Lyu, M.-S., Yang, Y.-L., Lin, E. P., Dunbrack, R., Birren, B., Cunningham, J., & Hunter, K. (1999). *Ier5*,a Novel Member of the Slow-Kinetics Immediate-Early Genes. Genomics, 55(3), 327–334. 10.1006/geno.1998.5679

Winkles, J. A. (1997). Serum- and Polypeptide Growth Factor-Inducible Gene Expression in Mouse Fibroblasts. In K. Moldave (Ed.), Progress in Nucleic Acid Research and Molecular Biology (Vol. 58, pp. 41–78). Academic Press. 10.1016/S0079-6603(08)60033-1

Winston, J. T., & Pledger, W. J. (1993). Growth factor regulation of cyclin D1 mRNA expression through protein synthesis-dependent and -independent mechanisms. Molecular Biology of the Cell, 4(11), 1133–1144. 10.1091/mbc.4.11.1133

Yamamoto, K. R., & Alberts, B. M. (1976). Steroid Receptors: Elements for Modulation of Eukaryotic Transcription. Annual Review of Biochemistry, 45(Volume 45, 1976), 721–746. 10.1146/annurev.bi.45.070176.003445

Yan, J., Chen, S.-A. A., Local, A., Liu, T., Qiu, Y., Dorighi, K. M., Preissl, S., Rivera, C. M., Wang, C., Ye, Z., Ge, K., Hu, M., Wysocka, J., & Ren, B. (2018). Histone H3 lysine 4 monomethylation modulates long-range chromatin interactions at enhancers. Cell Research, 28(2), 204–220. 10.1038/cr.2018.1

Zabidi, M. A., Arnold, C. D., Schernhuber, K., Pagani, M., Rath, M., Frank, O., & Stark, A. (2015). Enhancer–core-promoter specificity separates developmental and housekeeping gene regulation. Nature, 518(7540), 556–559. 10.1038/nature13994

Zambrano, S., Loffreda, A., Carelli, E., Stefanelli, G., Colombo, F., Bertrand, E., Tacchetti, C., Agresti, A., Bianchi, M. E., Molina, N., & Mazza, D. (2020). First Responders Shape a Prompt and Sharp NF-κB-Mediated Transcriptional Response to TNF-α. iScience, 23(9), 101529. 10.1016/j.isci.2020.101529

Zenklusen, D., Larson, D. R., & Singer, R. H. (2008). Single-RNA counting reveals alternative modes of gene expression in yeast. Nature Structural & Molecular Biology, 15(12), 1263–1271. 10.1038/nsmb.1514

Zhang, F., Biswas, M., Massah, S., Lee, J., Lingadahalli, S., Wong, S., Wells, C., Foo, J., Khan, N., Morin, H., Saxena, N., Kung, S. H. Y., Sun, B., Parra Nuñez, A. K., Sanchez, C., Chan, N., Ung, L., Altıntaş, U. B., Bui, J. M., … Lallous, N. (2023). Dynamic phase separation of the androgen receptor and its coactivators key to regulate gene expression. Nucleic Acids Research, 51(1), 99–116. 10.1093/nar/gkac1158

Zhou, T., & Feng, Q. (2022). Androgen receptor signaling and spatial chromatin organization in castration-resistant prostate cancer. Frontiers in Medicine, 9. https://www.frontiersin.org/articles/10.3389/fmed.2022.924087

