## Supplementary material for "Ligand-dependent Enhancer Activation Indirectly Modulates Non-target Promoters in a Chromatin Domain": Fig 1- Figure Supp 1

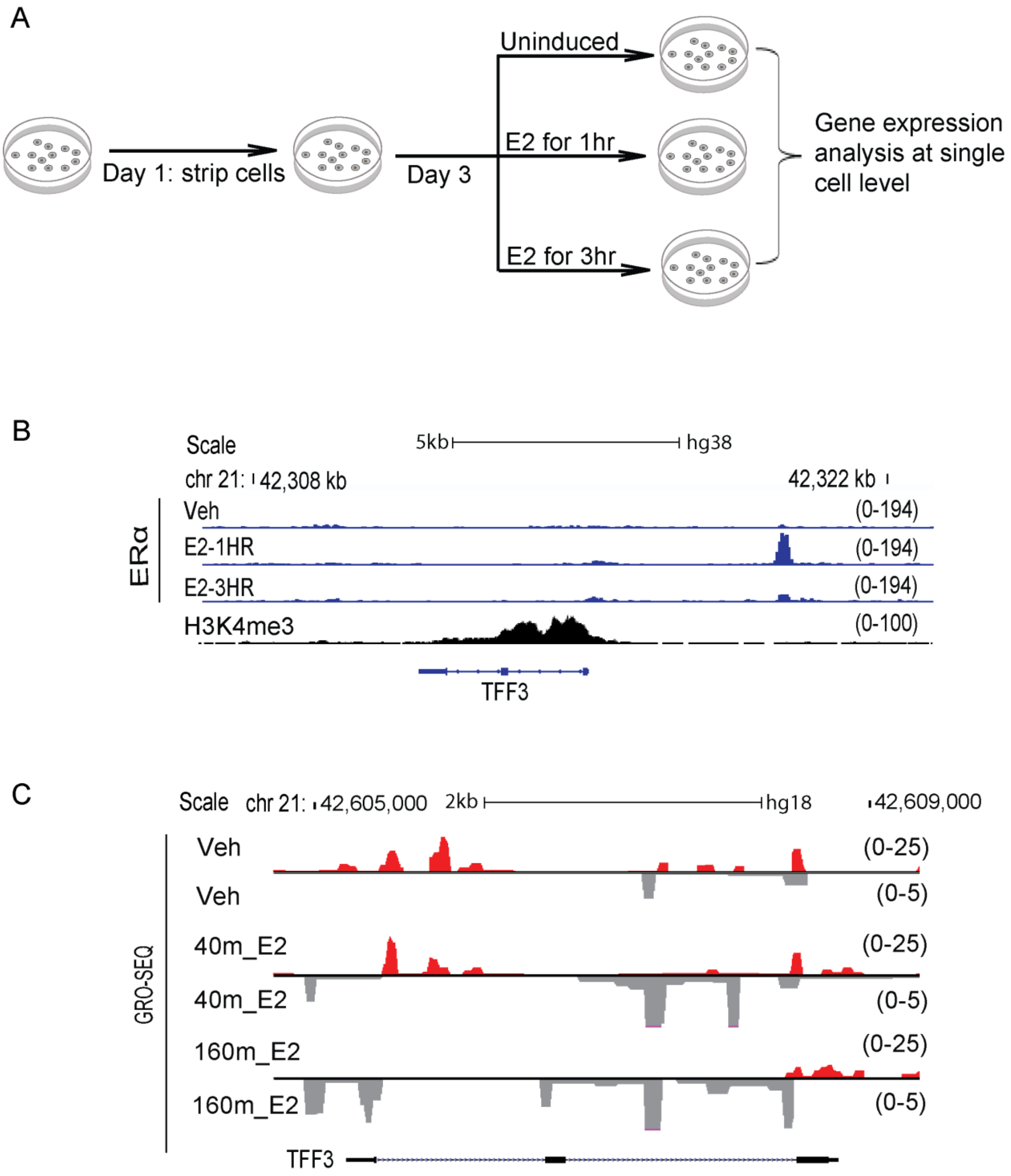


Figure 1—figure supplement 1

**Figure 1—figure supplement 1. Experimental design**

A. Schematic depicting the experimental design. Cells were cultured in complete media for 24h followed by stripping for 3 days. Finally, cells were induced for different durations with E2/Vehicle followed by different assays like smFISH, 4C-seq, smFISH-IF, etc.
B. Zoomed in region around TFF3 gene, UCSC genome browser snapshots showing the binding of ERα and H3k4me3 signal. First, second and third ERα ChIP-seq tracks are in vehicle treated , E2 treated for 1h and E2 treated for 3h in WT cells respectively
C. UCSC genome browser snapshots showing Gro-seq signal for *TFF3* gene. First, second and third Gro-seq tracks are from vehicle-treated, E2-40m and E2-160m in WT cells, respectively.
