## Supplementary material for "Ligand-dependent Enhancer Activation Indirectly Modulates Non-target Promoters in a Chromatin Domain": Fig 2- Figure Supp 1

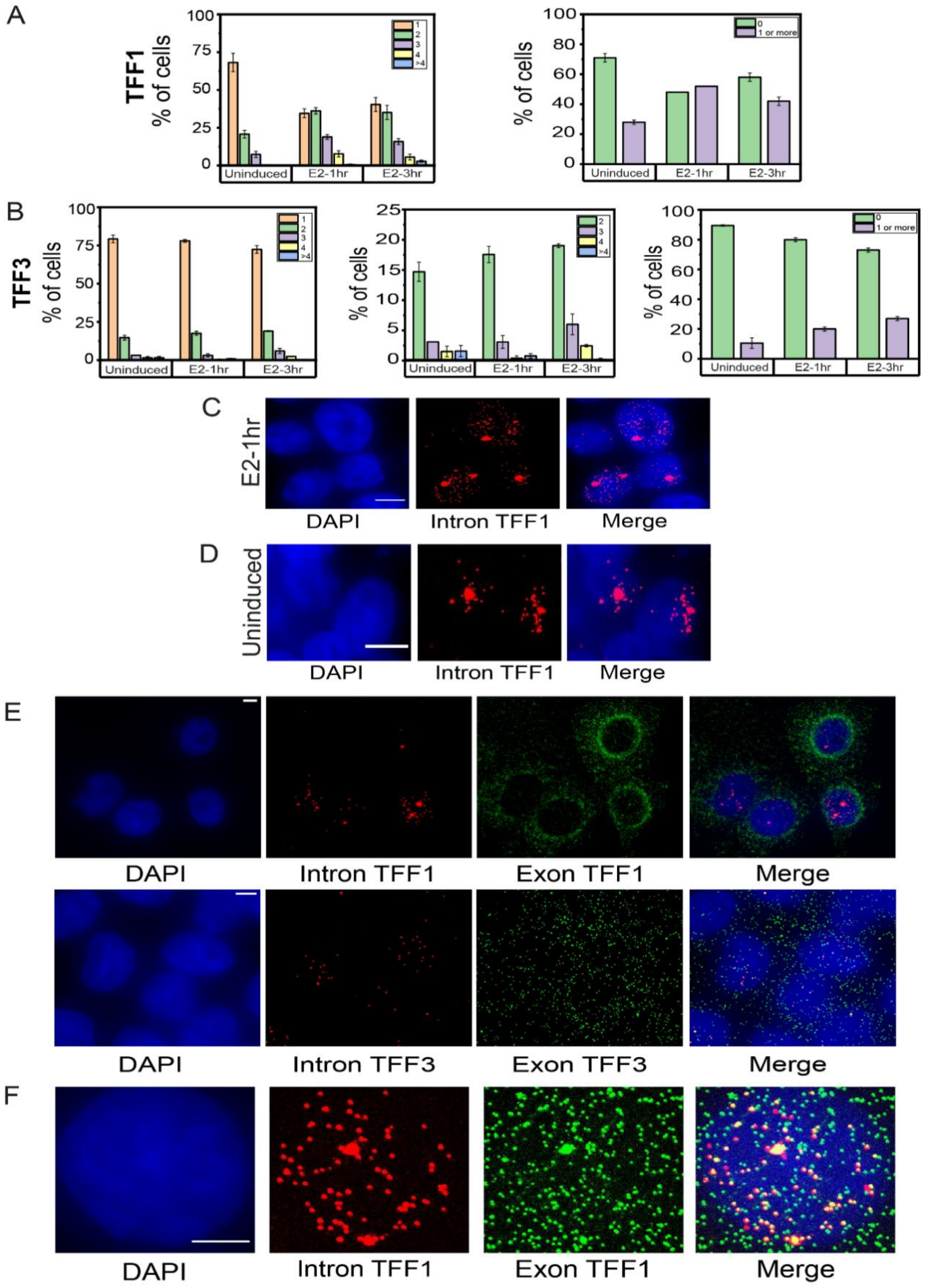


Figure 2—figure supplement 1

**Figure 2—figure supplement 1. Dynamic induction and RNA localization of *TFF1* and *TFF3* transcription across cell populations using smRNA FISH**

A. Bar graph depicting the percentage of cells with 1,2,3,4, or greater than 4 sites of transcription for TFF1 (left) is shown. The graph shows the mean of means from different repeats of the experiment, and error bars denote SEM (n>200, N=3). Only the cells with at least one allele firing were counted and cells with no alleles were not included in this. The graph on right shows the number of cells with zero or non-zero number of alleles firing.

B. Bar graph depicting the percentage of cells with 1,2,3,4 or greater than 4 sites of transcription for TFF3 (left) is shown. The graph shows the mean of means from different repeats of the experiment, and error bars denote SEM (n>200, N=3). Only the cells with at least one allele firing were counted and cells with no alleles were not included in this. The graph in the middle shows the number of cells with 2,3,4 or greater than 4 sites of transcription for TFF3.The graph on the right shows the number of cells with zero or non-zero number of alleles firing.

C. Images from single molecule RNA FISH experiment showing transcripts for InTFF1 in cells induced for 1 hour with E2. The image shows that when a single allele of TFF1 is firing, the transcripts show a more spatially restricted localisation. The scale bar is 5 microns.

D. Images from single molecule RNA FISH experiment showing transcripts for InTFF1 in uninduced cells. The image shows that when a single allele of TFF1 is firing and transcription is low, the transcripts show a more spatially restricted localisation. The scale bar is 5 microns.

E. 60X Representative images from a single molecule RNA FISH experiment showing transcripts for InTFF1 and ExTFF1 (top) and InTFF3 and ExTFF3 (bottom). The image shows that there is no intronic signal in the cytoplasm, while exonic signals can be found both in the nucleus and the cytoplasm. The scale bar is 5 microns.

F. 60X Representative images from single molecule RNA FISH experiment showing transcripts for InTFF1 and ExTFF1. The image shows that all intronic signals are colocalized with exonic signals, but all exonic signals are expectedly not colocalized with intronic signals, representing more mature mRNA. The scale bar is 5 microns.
