## Supplementary material for "Ligand-dependent Enhancer Activation Indirectly Modulates Non-target Promoters in a Chromatin Domain": Fig 2- Figure Supp 2

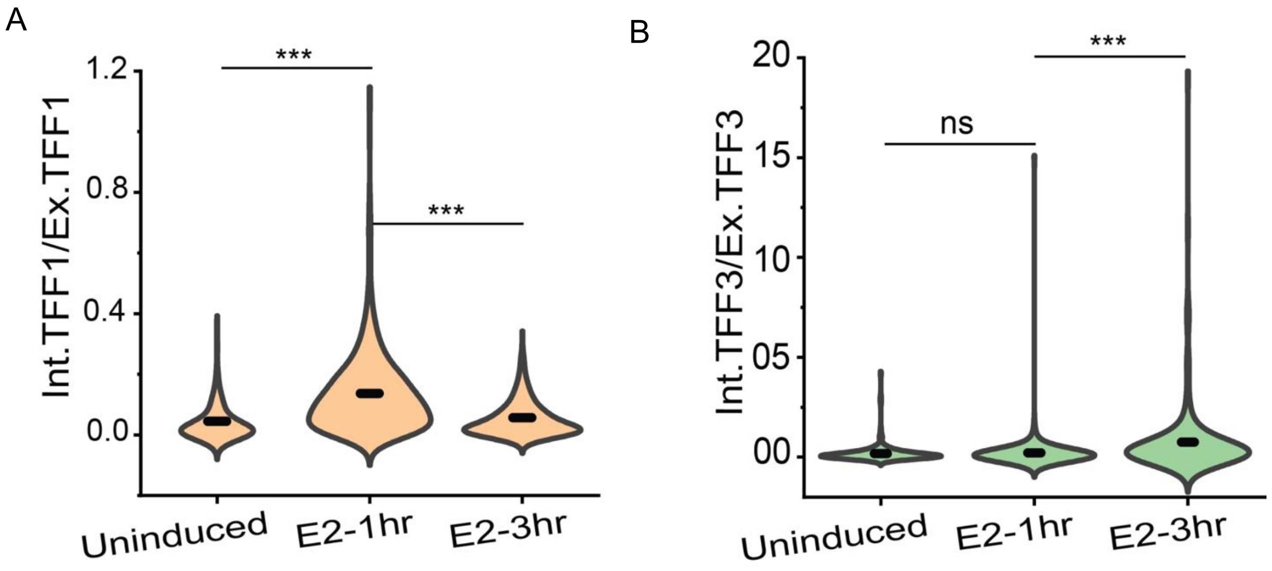


Figure 2—figure supplement 2

**Figure 2—figure supplement 2. *TFF1* and *TFF3* expressions show opposite trends during the E2-signaling time-course**

A. Violin plot showing the ratio of total intronic to absolute exonic *TFF1* counts are depicted. Absolute exonic counts are calculated by subtracting total intronic transcripts from total exonic transcripts. The graph shows the distribution of ratios combined from three different repeats. Error bars denote SEM. p-values were calculated by the Mann-Whitney test, and the significance is represented as: *** denotes $p < 0.001$, ** denotes $p < 0.01$, * denotes $p < 0.05$, and ns denotes $p > 0.05$.

B. Violin plot showing the ratio of total intronic to absolute exonic *TFF3* counts are depicted. Absolute exonic counts are calculated by subtracting total intronic transcripts from total exonic transcripts. The graph shows the distribution of ratios combined from three different repeats. Error bars denote SEM. p-values were calculated by the Mann-Whitney test, and the significance is represented as: *** denotes $p < 0.001$, ** denotes $p < 0.01$, * denotes $p < 0.05$, and ns denotes $p > 0.05$.
