## Supplementary material for "Ligand-dependent Enhancer Activation Indirectly Modulates Non-target Promoters in a Chromatin Domain": Fig 3- Figure Supp 1

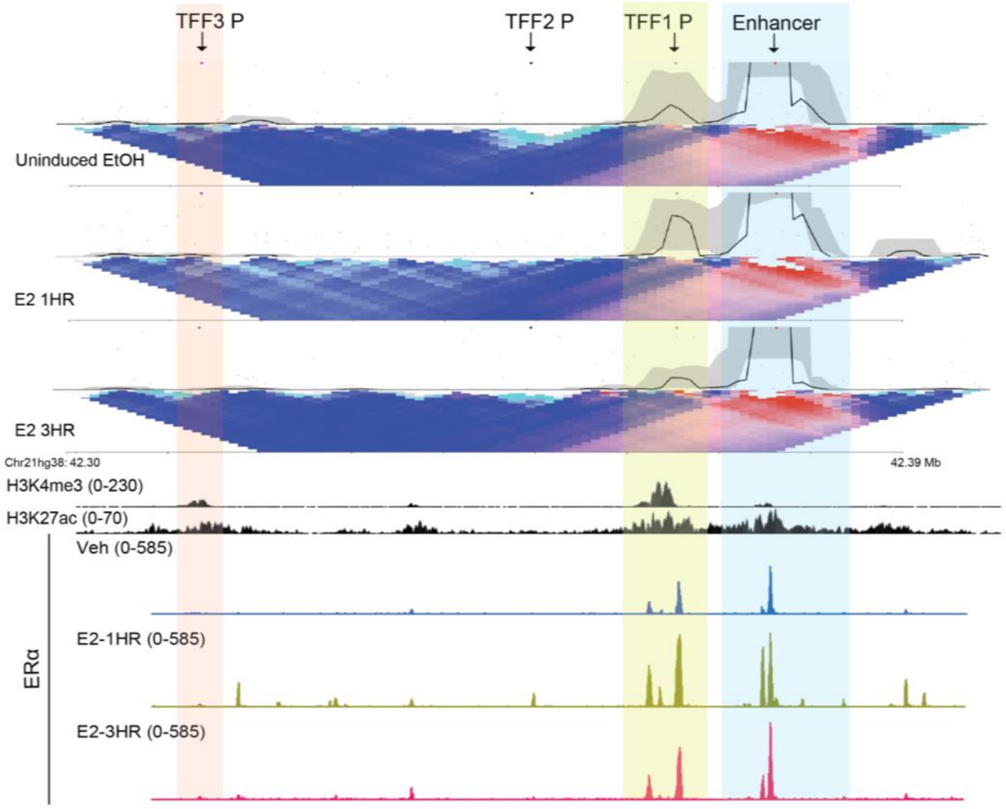


Figure 3—figure supplement 1

**Figure 3—figure supplement 1. *TFF1* enhancer does not change target promoters during signalling time-course**

4C-seq plot at *TFF1* enhancer viewpoint, the interaction with the promoter is highlighted in yellow. The plot is overlaid with H3K27ac, ERα ChIP signal, and gene annotations. There is no substantial interaction between the enhancer and *TFF3* locus at any time point, while the interaction between the enhancer and *TFF1* locus increases at 1hr and decreases at 3hr.
