## Supplementary material for "Ligand-dependent Enhancer Activation Indirectly Modulates Non-target Promoters in a Chromatin Domain": Fig 4- Figure Supp 1

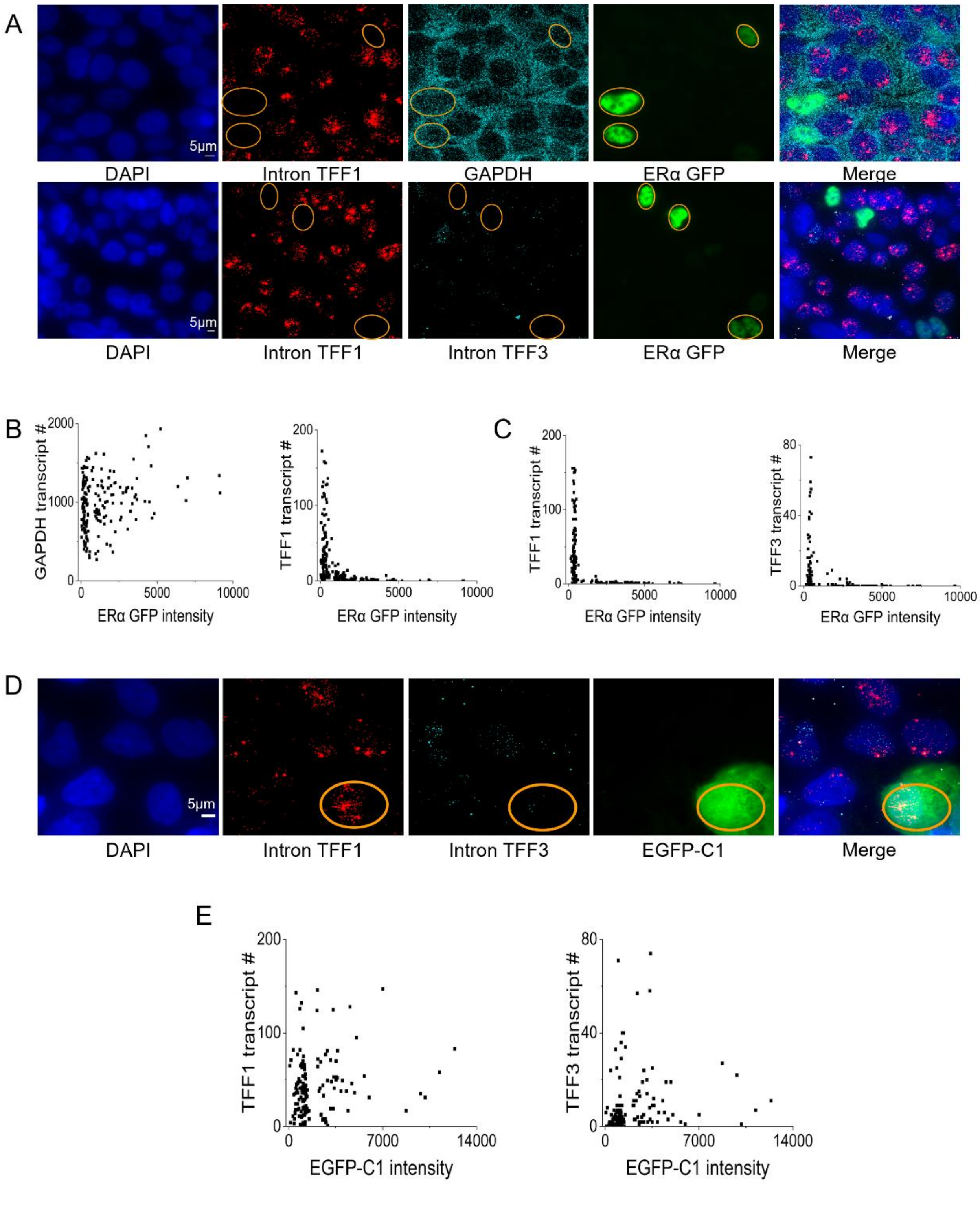


Figure 4—figure supplement 1

**Figure 4—figure supplement 1. Levels of ERα in the nucleus dictate the extent of *TFF1* and *TFF3* inductions**

A. Representative images from smFISH experiments showing In*TFF1* and GAPDH (top panel) and In*TFF1* and In*TFF3* (bottom panel) in cells overexpressing ERα-GFP. Yellow circles show cells that are GFP positive, and the transcripts associated with them. Note the visibly fewer *TFF1*, and *TFF3* transcripts in the ERα-GFP positive cells, while GAPDH transcripts remain indistinguishable. This shows that ERα-GFP overexpression specifically affects the transcription of E2-regulated genes like *TFF1* and *TFF3* and not a housekeeping gene like GAPDH. This is further quantified in B and C.

B. Scatter plots showing the distribution of ERα-GFP intensities with GAPDH or In*TFF1* transcript counts on a cell-by-cell basis from the experiment in the first row of A. Cells with high ERα-GFP expression can have high GAPDH expression, but such cells necessarily have low *TFF1* counts.

C. Scatter plots showing the distribution of ERα GFP intensities with In*TFF1* or In*TFF3* counts on a cell-by-cell basis from the experiment in the second row of A. As in B, high ERα-GFP expression leads to low expression of E2-regulated genes like *TFF1* or *TFF3*.

D. Representative images showing smFISH for In*TFF1* and In*TFF3* in cells overexpressing EGFP-C1. Yellow circles show cells that are GFP positive, and the transcripts associated with them. Just expressing EGFP in the same backbone as the ERα-GFP plasmid, does not downregulate *TFF1*, and *TFF3* expression; indicating that the downregulation effect is specific to ERα overexpression and not a generic effect of cell transfection. This is quantified in E.

E. Scatter plots showing the distribution of EGFP-C1 intensities with In*TFF1* or In*TFF3* transcript counts on a cell-by-cell basis. There is no obvious correlation between the levels of EGFP-C1 and *TFF1* or *TFF3*, and a cell with high EGFP levels may well have large *TFF1* or *TFF3* transcript counts.
