## Supplementary material for "Ligand-dependent Enhancer Activation Indirectly Modulates Non-target Promoters in a Chromatin Domain": Supp Tables

### Supplementary File 1a- InTFF3 probe sequence

| Probe Sequence (5' to 3' | Probe sequence name |
|---|---|
| catctttcctttttctttgt | Int_TFF3_1 |
| ctgaagtcagggtttttctc | Int_TFF3_2 |
| taaatgaaccttgtgcctgc | Int_TFF3_3 |
| ggcagggttgctatgcaatg | Int_TFF3_4 |
| acttgggttaatggagcagc | Int_TFF3_5 |
| ataaaagccatgccctcatt | Int_TFF3_6 |
| tttgaccctgctcagaagac | Int_TFF3_7 |
| cgtaattgcttatcctcctg | Int_TFF3_8 |
| tcttacattcccttctcttg | Int_TFF3_9 |
| ggaaaatggcaatcacgtgc | Int_TFF3_10 |
| cagaagttcactgcgtttca | Int_TFF3_11 |
| caagcctctatgtctgtatg | Int_TFF3_12 |
| ccccaatattttcatgtttc | Int_TFF3_13 |
| ctgaattgttgagcatgtcc | Int_TFF3_14 |
| tgcttttcactttttctgc | Int_TFF3_15 |
| cgccatagagatcattgcaa | Int_TFF3_16 |
| aaaccgtgacaggagaggca | Int_TFF3_17 |
| cttgcccctttgttttaaat | Int_TFF3_18 |
| ccttcattgtttgaccaaca | Int_TFF3_19 |
| cccttgaacccagaaacaaa | Int_TFF3_20 |
| cttctcgggaagtcacatac | Int_TFF3_21 |
| catttgcctaagtgtctttc | Int_TFF3_22 |
| gtctgttaaatgctcagctc | Int_TFF3_23 |
| ccttaggaagatgcactttc | Int_TFF3_24 |
| cgagttcaaccactgctgaa | Int_TFF3_25 |
| agtgtgtttgcttcactttg | Int_TFF3_26 |
| agcccagaaaggacagggag | Int_TFF3_27 |
| ttggaacaggtgtgtgtgtg | Int_TFF3_28 |
| taagtgtgaagcctctgctc | Int_TFF3_29 |
| aggggacagaagaggacagc | Int_TFF3_30 |
| tgcgatgccatcattttgaa | Int_TFF3_31 |
| gttactaaatcagttccctc | Int_TFF3_32 |
| gtttccacttctctcaagaa | Int_TFF3_33 |
| ggccgatttgcaaaaacagc | Int_TFF3_34 |
| ggggatgccacaaatgacaa | Int_TFF3_35 |
| cagagagagggagaggagat | Int_TFF3_36 |
| cacgatgtggggcagctaag | Int_TFF3_37 |
| agatggcttatagccaatcg | Int_TFF3_38 |
| tgagctggcacttggacatg | Int_TFF3_39 |
| tggatgctggttccaggaag | Int_TFF3_40 |
| cctaggacaaagggcaagag | Int_TFF3_41 |
| catcattgatcagggagccg | Int_TFF3_42 |

| | |
|---|---|
| agtggagctgaaggaacagg | Int_TFF3_43 |
| cactcagaggagagtgtagt | Int_TFF3_44 |
| cagggatcaggcagataaca | Int_TFF3_45 |
| tcgggttgctaagcaagata | Int_TFF3_46 |
| atgggggaatagccaagatt | Int_TFF3_47 |
| acagagcaaaggcttgtgtg | Int_TFF3_48 |

### Supplementary File 1b- ExTFF3 probe sequence

| Probe Sequence (5' to 3') | Probe sequence name |
|---|---|
| aaggggagaggtcagttca | Exn_TFF3_1 |
| ccagacagtttttctctcc | Exn_TFF3_2 |
| ctgcatgcctttgtcaagc | Exn_TFF3_3 |
| ttgtttgcttggggaaggc | Exn_TFF3_4 |
| tgtttgcacagctgctctg | Exn_TFF3_5 |
| aggaggcctcatttatgca | Exn_TFF3_6 |
| ctcaggactcgcttcatgg | Exn_TFF3_7 |
| cagcatgcagagcgctctg | Exn_TFF3_8 |
| aggacagcaaggccaggac | Exn_TFF3_9 |
| cacgtactcctcagcagag | Exn_TFF3_10 |
| acggcacactggtttgcag | Exn_TFF3_11 |
| actccttgggggtgacatg | Exn_TFF3_12 |
| cagggatcctggagtcaaa | Exn_TFF3_13 |
| ggggcttgaaacaccaagg | Exn_TFF3_14 |
| gtgcctcagaaggtgcatt | Exn_TFF3_15 |
| caatcacagccgggcaagg | Exn_TFF3_16 |
| gctgagatgaacagtgcct | Exn_TFF3_17 |
| cgggagcaaagggacagaa | Exn_TFF3_18 |
| actttcagcagaagcgctt | Exn_TFF3_19 |
| gacatcaggctccagatat | Exn_TFF3_20 |
| catgggacctttattcgtt | Exn_TFF3_21 |
| gaagaactgtcctcgggtg | Exn_TFF3_22 |
| acctcagaaagtctcaggc | Exn_TFF3_23 |
| ccacgacgcagcagaaata | Exn_TFF3_24 |
| cttagggagtcttttcctg | Exn_TFF3_25 |
| atccttgcatgcactgcag | Exn_TFF3_26 |
| gtgccagtctggattcaaa | Exn_TFF3_27 |
| ctaggctttcctgtgacgt | Exn_TFF3_28 |
| gaagcggcacttacagtgt | Exn_TFF3_29 |
| gtatttttctgctttccc | Exn_TFF3_30 |
| ttattggcttgctgtcttc | Exn_TFF3_31 |
| ggaccactttggaagacag | Exn_TFF3_32 |
| gtagcgagagtggttgtga | Exn_TFF3_33 |
| acccaggaatagtacaagt | Exn_TFF3_34 |

| | |
|---|---|
| gcctgtttgttcataatgt | Exn_TFF3_35 |

### Supplementary File 1c- InTFF1 probe sequence

| Probe Sequence (5' to 3') | Probe sequence name |
|---|---|
| acagagcaggaagaagcacg | Int_TFF1_1 |
| tgacacttgggaggattgta | Int_TFF1_2 |
| ccaacacttcctctttgaaa | Int_TFF1_3 |
| aaagcactattctgagaccc | Int_TFF1_4 |
| gggagatgttggcatgaaca | Int_TFF1_5 |
| catttgggcctatctggatg | Int_TFF1_6 |
| caggaacctgtaggagaggg | Int_TFF1_7 |
| atgagagaggtggctttgac | Int_TFF1_8 |
| cagggaatatgggttgccag | Int_TFF1_9 |
| caaccgatccacttgacaac | Int_TFF1_10 |
| ttgagatgcaaacacttccc | Int_TFF1_11 |
| atccattctgcaggtaaagg | Int_TFF1_12 |
| cacacagcatttcctgacag | Int_TFF1_13 |
| aaagtgcaagtcgcagatgc | Int_TFF1_14 |
| aataagttattcagctcccc | Int_TFF1_15 |
| ccaccttgaaactgtaccta | Int_TFF1_16 |
| aacgtagaaagccctttctc | Int_TFF1_17 |
| ctctggaaacccttgctttg | Int_TFF1_18 |
| ccacagcagagatcaagagg | Int_TFF1_19 |
| agttcgttctgtacaccgag | Int_TFF1_20 |
| aaagacgctctgagccttga | Int_TFF1_21 |
| gtaacaggatcatggacctg | Int_TFF1_22 |
| ttccacacacgaatgcacta | Int_TFF1_23 |
| ctttgtctttgtcgcatgat | Int_TFF1_24 |
| tcagggaggaagatttccag | Int_TFF1_25 |
| tggtgagggaggatcatcat | Int_TFF1_26 |
| cactgtgggggtcaagagag | Int_TFF1_27 |
| cgacgtgaaggtgatcatcg | Int_TFF1_28 |
| tgggaaggatccgtgttcag | Int_TFF1_29 |
| gtctttggagaaagtgttcc | Int_TFF1_30 |
| cttgtctgctttgctatcag | Int_TFF1_31 |
| ataggaggggaggaactcag | Int_TFF1_32 |
| agcagggggtgaaagaggag | Int_TFF1_33 |
| taggatctggtggttgacag | Int_TFF1_34 |
| cgcggaatcaaaggtctcag | Int_TFF1_35 |
| tctaacaccagagcttcagg | Int_TFF1_36 |
| cgcaggatcttatgcaagag | Int_TFF1_37 |
| agaggaggaagacacgtgga | Int_TFF1_38 |
| aagcagggcagtgaggaaag | Int_TFF1_39 |
| cagagggtacaagctgtgaa | Int_TFF1_40 |

| | |
|---|---|
| attgcaggtaaggtgagagg | Int_TFF1_41 |
| gtgtgttcaggaggtggaaa | Int_TFF1_42 |
| ttcaccaggtgaagatggag | Int_TFF1_43 |
| aggttgaacaaagcaggggg | Int_TFF1_44 |
| atgtgcgagacaacgcaagg | Int_TFF1_45 |
| tctgatggaggagaaaggca | Int_TFF1_46 |
| ttgtggactttgcatctttc | Int_TFF1_47 |
| gctcagggaaaatgcaaggt | Int_TFF1_48 |

### Supplementary File 1d- ExTFF1 probe sequence

| Probe Sequence (5' to 3') | Probe sequence name |
|---|---|
| gcgaccccgagtcagggat | Exn_TFF1_1 |
| tgcctcctctctgctccaa | Exn_TFF1_2 |
| cttgttctccatggtggcc | Exn_TFF1_3 |
| caggaccagggcgcagatc | Exn_TFF1_4 |
| gccgagggccagcatggac | Exn_TFF1_5 |
| tgtctgggcctcggccagg | Exn_TFF1_6 |
| ggggccactgtacacgtct | Exn_TFF1_7 |
| ccacaattctgtctttcac | Exn_TFF1_8 |
| gagggcgtgacaccaggaa | Exn_TFF1_9 |
| cagcccttatttgcacact | Exn_TFF1_10 |
| cacgaacggtgtcgtcgaa | Exn_TFF1_11 |
| gatagaagcaccaggggac | Exn_TFF1_12 |
| gagggacgtcgatggtatt | Exn_TFF1_13 |
| tcgtataaaaaggccatac | Exn_TFF1_14 |
| attctaaatcttcagaacc | Exn_TFF1_15 |
| gtcttaaatgacttttcta | Exn_TFF1_16 |
| atgctgatcagagcctctg | Exn_TFF1_17 |
| tgtgtaaaggcatagctgg | Exn_TFF1_18 |
| caccactggcggccgtgac | Exn_TFF1_19 |
| gactcaggctaccccattg | Exn_TFF1_20 |
| ttcctggacctgaatgcag | Exn_TFF1_21 |
| tcccttagccctgcctttc | Exn_TFF1_22 |
| gtctaaaattcacactcct | Exn_TFF1_23 |
| caggcagatccctgcagaa | Exn_TFF1_24 |
| ggacggcaccgcgtcagga | Exn_TFF1_25 |
| tgggactaatcaccgtgct | Exn_TFF1_26 |
| ggtggaggtggcagccgag | Exn_TFF1_27 |
| agaagcgtgtctgaggtgt | Exn_TFF1_28 |
| tgtgagccgaggcacagct | Exn_TFF1_29 |
| gtcagagcagtcaatctgt | Exn_TFF1_30 |
| ggccaattttgagtagtca | Exn_TFF1_31 |
| atcgatctcttttaatttt | Exn_TFF1_32 |

**Supplementary File 1e- 4C Seq oligo sequences**

| Sequence name | Sequence (5' to 3') |
|---|---|
| 4C TFF1 Enhancer barcode1 F | AATGATACGGCGACCACCGAGATCTACACTCTTTCCCTACACGACGCTCTTCCGATCTAGAAGTCCTTGAGTGGGGAGATGATC |
| 4C TFF1 Enhancer barcode2 F | AATGATACGGCGACCACCGAGATCTACACTCTTTCCCTACACGACGCTCTTCCGATCTAGCGACCCTTGAGTGGGGAGATGATC |
| 4C TFF1 Enhancer barcode3 F | AATGATACGGCGACCACCGAGATCTACACTCTTTCCCTACACGACGCTCTTCCGATCTCCAATTCCTTGAGTGGGGAGATGATC |
| 4C TFF1 R | CAAGCAGAAGACGGCATACGAGATAGGTCGCAGTGACTGGAGTTCAGACGTGTGCTCTTCCGATCTCCGCTGTTGCGACACACCAGAGAGC |

**Supplementary File 1f- qPCR oligo sequences**

| Sequence name | Sequence (5' to 3') |
|---|---|
| TFF1 US F | GCCCTTATTTGCACACTGGG |
| TFF1 US R | CATCAAGGAATTCAGCCCAC |
| TFF3 US F | CAATCACAGCCGGGCAAGGGT |
| TFF3 US R | GGGAGGAGGCAGCACTAGGT |
| GAPDH F | CGCTCTCTGCTCCTCCTGTT |
| GAPDH R | CCATGGTGTCTGAGCGATGT |
